# Molecular Insights into the Recognition of Acetylated Histone Modifications by the BRPF2 Bromodomain

**DOI:** 10.1101/2022.02.20.481182

**Authors:** Soumen Barman, Anirban Roy, Jyotirmayee Padhan, Babu Sudhamalla

## Abstract

HBO1 (HAT bound to ORC), a member of the MYST family of histone acetyltransferases (HATs), was initially identified as a binding partner of the origin recognition complex (ORC) that acetylates free histone H3, H4, and nucleosomal H3. It functions as a quaternary complex with the BRPF (BRPF1/2/3) scaffolding protein and two accessory proteins, ING4/5 and Eaf6. BRPF2 interaction with HBO1 has been shown to be important for regulating H3K14 acetylation during embryonic development. However, how the BRPF2 directs the HBO1 HAT complex to chromatin to regulate its HAT activity towards nucleosomal substrates remains unclear. Our findings reveal novel interacting partners of the BRPF2 bromodomain that recognizes different acetyllysine residues on the N-terminus of histone H4, H3, and H2A and preferentially binds to H4K5ac, H4K8ac, and H4K5acK12ac modifications. Further, mutational analysis of BRPF2 bromodomain coupled with ITC binding and pull-down assays on the histone substrates identified critical residues responsible for acetyllysine binding. Moreover, the BRPF2 bromodomain could enrich H4K5ac mark-bearing mononucleosomes compared to other acetylated H4 marks. Consistent with this, ChIP-seq analysis revealed that BRPF2 strongly co-localizes with HBO1 at histone H4K5ac and H4K8ac marks near the TSS in the genome. Together, our study provides novel insights into how the histone binding function of the BRPF2 bromodomain directs the recruitment of the HBO1 HAT complex to chromatin to regulate gene expression.

## INTRODUCTION

Histones are the basic proteins that package the DNA and are often subjected to a variety of post-translational modifications (PTMs), such as acetylation, methylation, phosphorylation, and ubiquitylation.^1–3^ These PTMs are reversible, and they are dynamically regulated by specialized class of proteins called ‘writers’ and ‘erasers’, which enzymatically add and remove the PTM marks, respectively.^4^ Different patterns of PTMs, which recruit the effector proteins to the chromatin, acts as an ‘off-on switch’ to regulate gene expression.^5^ Histone PTMs are recognized by another class of proteins called ‘readers’; these proteins bind to the specific PTM with the help of specialized domains and act as a scaffold for the recruitment of other regulatory complexes for downstream transcriptional regulation.^6^

Initially discovered 50 years ago, as a unique modification in histones, lysine acetylation is currently the most widely studied histone PTM.^5^ Acetylation marks are now known to also occur on thousands of nonhistone proteins in virtually every cellular compartment.^7–9^ This modification is recognized by specialized reader modules called bromodomains, which share a conserved globular fold and are composed of a left-handed bundle of four α-helices αZ, αA, αB, and αC.^10^ The helices are linked by two loop regions (ZA and BC), which form the hydrophobic acetyllysine (Kac) binding pocket.^10^ Moreover, the bromodomain and the acetyllysine binding is stabilized by hydrogen bonds with water molecules and a conserved asparagine ‘anchor’ residue in the bromodomain. Sixty-one different bromodomains have been identified so far, which recognize multiple acetylated lysine residues on the N-terminus of the histone tail.^10, 11^ Although the acetyllysine recognition occurs in the conserved structural fold of the bromodomain, selectivity is achieved through variation in the ZA and BC loop regions.^10^

The bromodomain and PHD finger (BRPF) family proteins are a sub-class of the bromodomain family and include BRPF1, BRPF2, and BRPF3.^10, 11^ These proteins contain an N-terminal PZP domain (two PHD fingers connected through a zinc knuckle), a bromodomain, and a C-terminal PWWP (Pro-Trp-Trp-Pro) domain (Figure 1A).^12^ BRPFs are chromatin binding scaffolding proteins and are components of the histone acetyltransferases (HATs) complexes.^13^ For example, BRPF1 forms a tetrameric complex with MOZ/MORF HAT, which acetylates several lysine residues on histone H3 and H4.^14–17^ Alternatively, BRPF2/3 associates with HBO1 HAT (Figure 1B), which acetylates K5, K8, and K12 of histone H4 and K14 of H3.^18–21^ BRPF family proteins stimulate and modulate the enzymatic activity of HATs and promote acetylation of histone lysine residues during transcription.^22–24^

**Figure 1.**
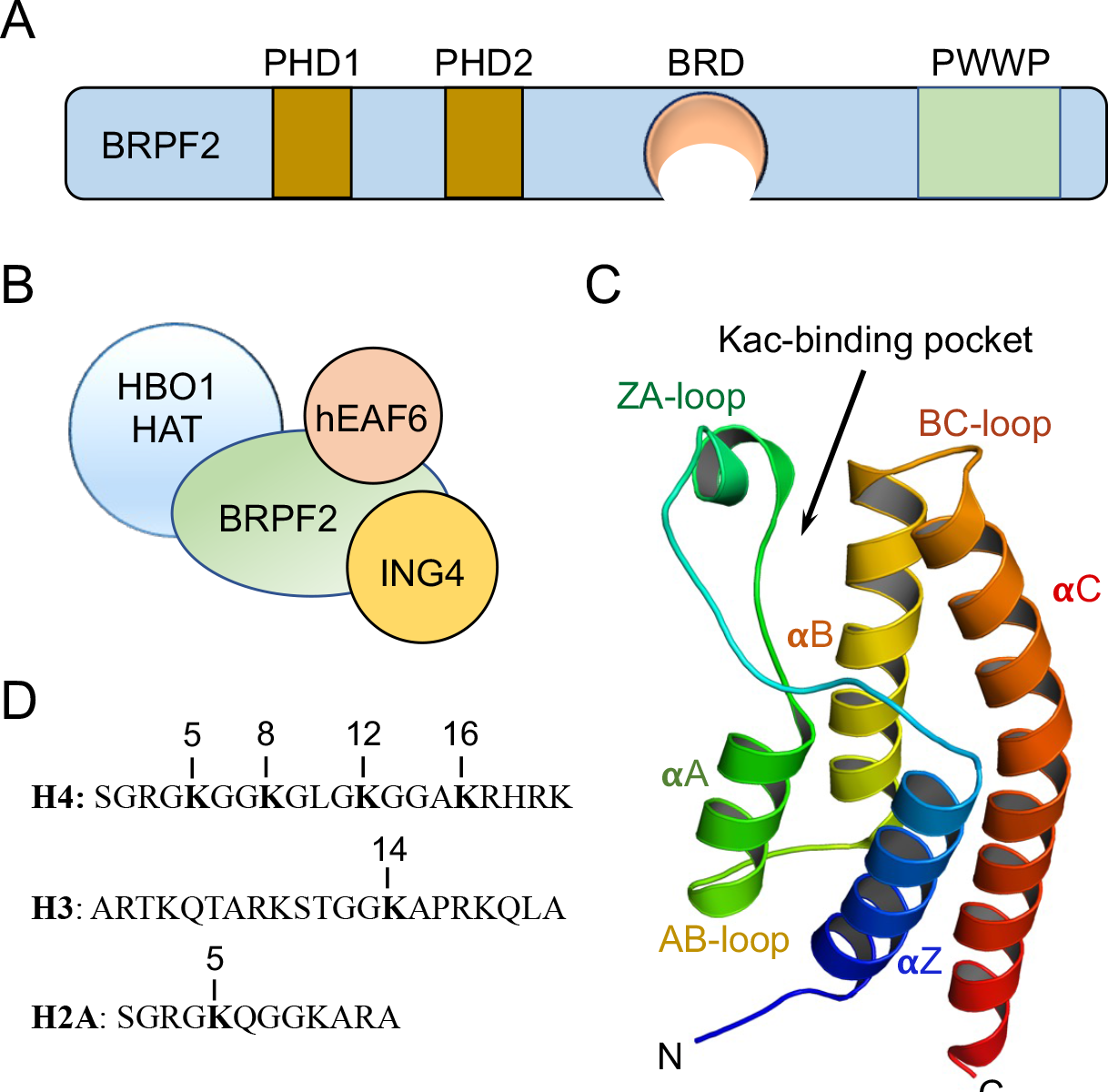
The BRPF2 bromodomain and histone ligands were characterized in this study. (A) Domain organization of the BRPF2 contains two plant homeodomain (PHD) fingers, one bromodomain (BRD), and one proline-tryptophan-tryptophan-proline (PWWP) domain. (B) BRPF2 forms a complex with HBO1 HAT and two other proteins. (C) Cartoon representation showing the crystal structure of the BRPF2 bromodomain (PDB: 3RCW). The secondary structure elements are labeled. (D) Histone H4, H3, and H2A peptide sequences and the highlighted lysine residues were subjected to acetylation.

The association of BRPFs with HATs plays a crucial role in development and disease progression. For example, the BRPF1-MOZ complex is essential for *HOX* gene expression in vertebrate segmentation and patterning.^25^ In both zebrafish and medaka fish models, BRPF1 mutation decreases *HOX* gene expression, which impairs proper skeletal development.^25, 26^ Similarly, the BRPF2-HBO1 complex regulates transcriptional activation of genes involved in the differentiation of erythrocytes in mice.^21^ Furthermore, BRPF2 deficient embryos develop severe anemia and do not survive due to impaired fetal liver erythropoiesis.^21^ The BRPF family proteins are also involved in bone degradation, and pharmacological inhibition of their bromodomain prevented osteoclastogenesis in murine and human cells.^27^ Apart from their role in development, BRPFs are misregulated in many diseases, including cancer. BRPF1 is among the most significantly upregulated gene in hepatocellular carcinoma.^28^ It increases the expression of critical oncogenes like *E2F2* and *EZH2* by facilitating their promoter acetylation in H3K14 through MOZ/MORF complex.^28^ BRPFs are increasingly becoming an attractive therapeutic target in cancer and various other diseases, attributed to their role in transcription and cell cycle.^28^ Potent and selective inhibitors for the BRPF family bromodomains have been developed in recent years. PFI-4 is highly selective for the BRPF1B isoform, BAY-299 is a potent and selective inhibitor for the BRPF2 bromodomain, and (OF-1, NI-57) have been reported as pan-BRPF bromodomain inhibitors.^29, 30^

The BRPF proteins recruit to chromatin by coordinating with their various domains that interact with histone and DNA.^24, 31^ The PZP domain binds unmethylated histone H3K4 while PWWP recognizes the H3K36me3 mark and the bromodomains of the BRPF family have a high affinity for multiple acetylated histone marks.^31–33^ Recent studies on the BRPF1 bromodomain revealed that it recognizes multiple acetyllysine residues on the histone H2A, H3, and H4 and preferentially binds to diacetylated histone H4K5acK8ac and H4K5acK12ac ligands.^34, 35^ Similarly, the BRPF3 bromodomain has a strong affinity for histone ligands such as H4K5ac and H4K5acK12ac.^36^ Although, it is reported that the BRPF2 bromodomain binds weakly to diacetylated histone H4K5acK8ac ligand,^37^ there is still limited histone ligand information available for the BRPF2 bromodomain (Figure 1C). We sought to identify novel acetylated histone ligands recognized by the BRPF2 bromodomain through a series of computational, biophysical, and biochemical methods. We report that the BRPF2 bromodomain binds to multiple acetyllysine residues on the N-terminal tails of histone H4, H3, and H2A. Moreover, we have characterized the mode of acetyllysine binding in the hydrophobic binding pocket of the BRPF2 bromodomain. And further validated the co-localization of BRPF2 with the identified histone targets in the genome.

## MATERIALS AND METHODS

### Peptide Synthesis

All histone peptides were synthesized by (GL Biochem Ltd., and S. Biochem Ltd.) and purified by HPLC to 98% purity. The integrity of the purified peptides were confirmed by LC-ESI mass spectrometry (Figure S9-S34 and Table S1). Peptide concentrations were determined based on the observation that 1mg/ml peptide generates an absorbance value (A_205_) of 30 at 205 nm.^38^

**Table 1.**
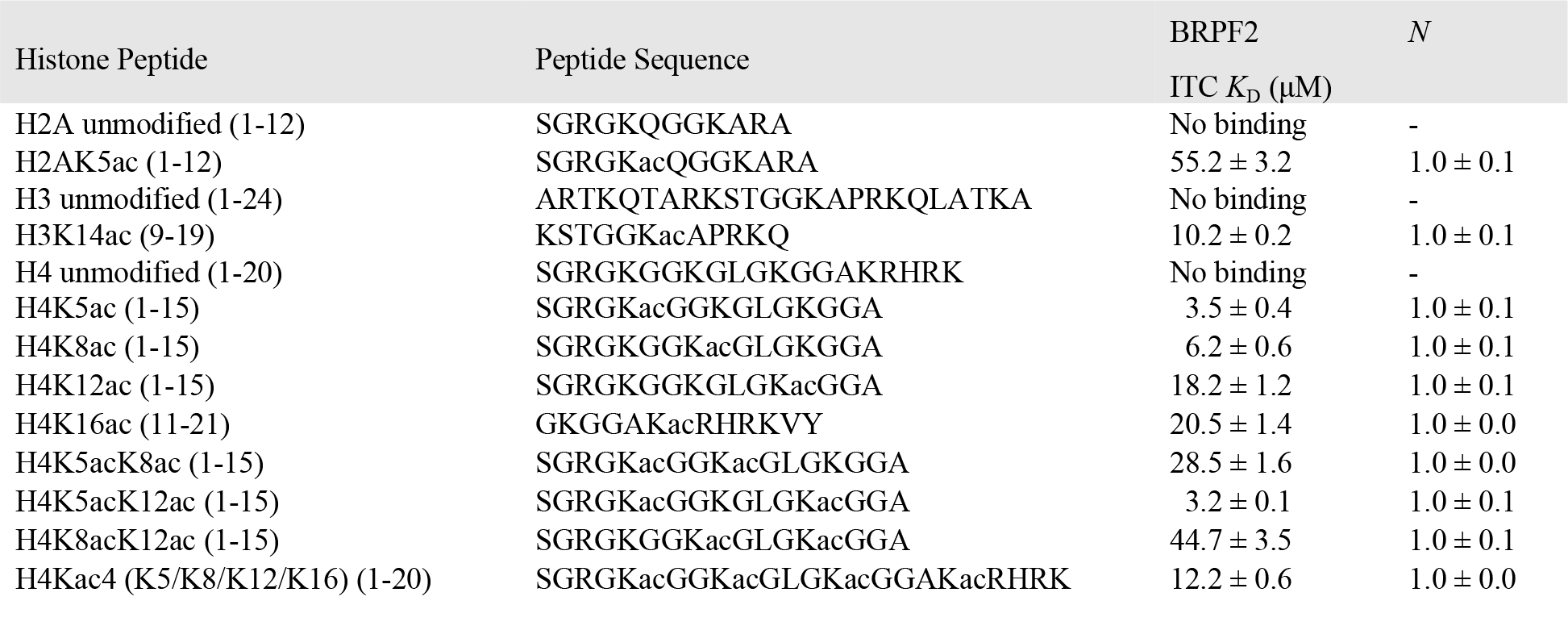
Dissociation constants of the interactions between BRPF2 bromodomain and acetylated histone peptides as measured by ITC

### Expression and Purification of BRPF2 Bromodomain

The protein expression and purification were done by methods described previously.^36, 39^ The N-terminal His_6_-tagged wild-type BRPF2 bromodomain plasmid (a gift from Nicola-Burgess-Brown, addgene ID: 39095) was transformed into One Shot BL21 star (DE3) *E. coli* competent cells (Invitrogen cat#C601003) using pNIC28- Bsa4 kanamycin-resistant vectors. A single colony was picked up and grown overnight at 37°C in 10 mL of Luria-Bertani (LB) broth in the presence of 50 μg mL^−1^ kanamycin. The culture was diluted 100-fold and allowed to grow at 37°C to an optical density (OD_600_) of 1.0. Protein expression was induced overnight at 17°C with 0.5 mM IPTG in an Innova 44 Incubator shaker (New Brunswick Scientific). Proteins were purified as follows: Harvested cells were resuspended in 15 mL lysis buffer (50 mM HEPES pH 7.5, 300 mM NaCl, 5 mM β-mercaptoethanol, 5% glycerol, 25 mM imidazole, Lysozyme, DNase, and 1:200 (v/v) Protease Inhibitor Cocktail III (Calbiochem). The cells were lysed by pulsed sonication and centrifuged at 13000 rpm for 40 min at 4°C. According to the manufacturer’s instructions, the soluble extracts were subject to Ni-NTA agarose resin (QIAGEN cat#30210). After passing 20 volumes of washing buffer (50 mM HEPES pH 7.5, 300 mM NaCl, 5 mM β-mercaptoethanol, 5% glycerol, and 25 mM imidazole), proteins were eluted with a buffer containing 50 mM HEPES pH 7.5, 300 mM NaCl, 5 mM β-mercaptoethanol, 5% glycerol, and 250 mM imidazole. Proteins were further purified by gel filtration chromatography (Superdex-75) using AKTA pure FPLC system (GE Healthcare) with a buffer containing 50 mM HEPES pH 7.5, 200 mM NaCl, and 5% glycerol. Purified proteins were concentrated using Amicon Ultra-10k centrifugal filter device (Merck Millipore Ltd.), and the concentration was determined using Bradford assay kit (Bio-Rad Laboratories) with BSA as a standard. The proteins were aliquoted and stored at -80°C before use.

Point mutations were introduced into the pNIC28-Bsa4 BRPF2 bromodomain construct to generate I586F, F587A, Y599F, N642A, and F648A mutants (Table S2) using the QuikChange Lightning site-directed mutagenesis kit following manufacturer’s protocol. The resulting mutant plasmids were confirmed by DNA sequencing. All the mutants were expressed and purified following the procedure used for wild-type BRPF2 bromodomain.

**Table 2.** Dissociation constants of the wild-type BRPF2 bromodomain and its mutants with histone H4K5ac (1-15) peptide as measured by ITC

| BRPF2 | H4K5ac (1-15) | *N* |
| --- | --- | --- |
| | ITC $K_D$ (μM) | |
| WT | $3.5 \pm 0.4$ | $1.0 \pm 0.1$ |
| I586F | $3.6 \pm 0.2$ | $1.0 \pm 0.0$ |
| F587A | $25.2 \pm 0.2$ | $1.0 \pm 0.1$ |
| Y599F | No binding | - |
| N642A | No binding | - |
| F648A | $45.2 \pm 0.3$ | $1.0 \pm 0.1$ |

### Isothermal Titration Calorimetry Measurements

ITC binding experiments were performed as previously described.^36, 39^ Briefly, experiments were carried out on a MicroCal PEAQ-ITC instrument (Malvern). Experiments were conducted at 15°C while stirring at 750 rpm. Buffers of protein and peptides were matched to 50 mM HEPES, pH 7.5, 200 mM NaCl, and 5% glycerol. Each titration comprised one initial injection of 0.4 μL lasting 0.8s, followed by 18 injections of 2 μL lasting 4s each at 2.5 min intervals. The initial injection was discarded during data analysis. The microsyringe (40 μL) was loaded with a peptide sample solution, and the peptide concentration varied from 1.5 mM to 2 mM. It was injected into the cell (200 μL), occupied by a protein concentration of 80 to 100 μM. All the data was fitted to a single binding site model using the MicroCal PEAQ-ITC analysis software to calculate the stoichiometry (N), the binding constant (*K*_D_), enthalpy (ΔH), and entropy (ΔS) of the interaction. The final titration figures were prepared using OriginPro 2020 software (OriginLab).

### Protein-Peptide Docking

Molecular docking was performed by following the method for docking bromodomain with acetylated histone peptide as previously described.^36, 39^ The crystal structure of the BRPF2 bromodomain (PDB: 3RCW) was used for molecular docking studies, and the polar hydrogen atoms were added using PyMoL software. The histone H4 coordinates were retrieved from the RCSB protein data bank, and the selected lysine residues were covalently modified using the PyTM plugin of PyMoL (version 1.8.4.0) software. The protein and histone peptide input structures were prepared, such as adding charges, assigning the atom types, and detecting the root for the ligand molecules were done by using the AutoDock tools. The grid box dimensions were set to 60 X 58 X 65 Å along the *X, Y*, and *Z* axes with a default grid spacing of 1Å, which covered the entire protein. Molecular docking of each modified histone peptide with BRPF2 bromodomain was performed using AutoDockVina 1.1.2.^40^ Molecular interaction between each peptide and bromodomain was analyzed using PyMOL (version 2.3.4) software package.

### Molecular Dynamics

The computational modeling and MD simulation were performed as previously described with slight modifications.^36, 39^ The best-ranked docked conformations of BRPF2 bromodomain and acetylated histone H4 peptides were subjected to 200 ns of MD simulations with GROMACS (version 2018.3) software package using the GROMOS96 54A7 force field.^41, 42^ In order to set up the MD simulations for the protein-peptide complexes, the topology parameters of BRPF2 bromodomain and acetylated histone H4 peptides were created using GROMACS and PRODRG2 server, respectively.^43^ The prepared protein-peptide complexes were then solvated in a dodecahedron box with a distance of 1.0 nm between the complex and edge of the solvated box. The solvated system was neutralized by adding sodium and chloride ions in the simulation. To ensure the complex’s steric clashes or geometry, the system was energy minimized using the LINCS constraints and steepest descent algorithms followed by system equilibration under NVT and NPT ensembles. Final MD simulations of bromodomain and peptide complexes were carried out at 300 K temperature, 1 atm pressure and 2 fs time step for 200 ns. The final MD trajectories were analyzed to calculate the RMSD (root mean square deviation), and RMSF (root mean square fluctuation) values by using the standard GROMACS functions.

### Mammalian Cell Culture

HeLa cells were grown in DMEM supplemented with 10% fetal bovine serum and antibiotics (Penicillin-Streptomycin cocktail) in a humidified atmosphere containing 5% CO_2_. Cells at ∼90% confluence stage were treated with 2 μM of histone deacetylase inhibitor Trichostatin A (TSA, cat# T8552, Sigma) dissolved in DMSO to generate hyperacetylated histones. Twenty hours post-treatment, media was removed and rinsed the cells with cold PBS buffer. The harvested cells were rewashed with cold PBS buffer and frozen as a dry pellet at -80 °C.

### Acid Extraction of Histones

Histones were extracted from HeLa cells using the acid-extraction protocol as previously described.^44^ Briefly, frozen cell pellets from 100 mm dish of HeLa cells were resuspended in 1 ml of hypotonic lysis buffer (10 mM Tris-HCl pH 8.0, 1 mM KCl, 1.5 mM MgCl_2_, 1 mM DTT, 1 mM PMSF, and protease inhibitor cocktail) and incubated for 30 min on nutator at 4°C. Samples were then centrifuged (10,000 x *g*, 10 min at 4°C), and the supernatant was discarded entirely. The nuclei resuspended in 800 μL of 0.4 N H_2_SO_4_, vortexed intermittently for 5 min, and further incubated at 4°C on a nutator for overnight. The nuclear debris was pelleted by centrifugation (16000 x *g*, 10 min at 4°C), and the supernatant containing histones were collected. The histones were precipitated by adding 264 μL TCA (Trichloroacetic acid) drop by drop to histone solution and inverted the tube several times to mix the solutions and incubated the samples on ice for 30 min. Finally, histones were pelleted by centrifugation (16000 x *g*, 10 min at 4°C), and the supernatant was discarded. The histones pellet was washed twice with ice-cold acetone, followed by centrifugation (16000 x *g*, 5 min at 4°C), and carefully removed the supernatant. The histone pellet was air-dried for 20 min at room temperature and subsequently dissolved in an appropriate volume of ddH2O and transferred into a fresh tube. The aliquoted histones were stored at -80°C until use.

### Isolation of Mononucleosomes From HeLa Cells

Nucleosomes were isolated from HeLa cells by following the protocol as previously described.^45, 46^ Briefly, the cells were harvested after twenty hours of post-treatment with 2 μM histone deacetylase inhibitor (TSA) and washed twice in PBS buffer. The cells were resuspended in buffer A (10 mM HEPES pH 7.9, 10 mM KCl, 1.5 mM MgCl_2_, 340 mM sucrose, 10% (v/v) glycerol, protease inhibitor cocktail (Pierce), 1 μg/mL TSA, 5 mM 2-mercaptoethanol), and an equivalent volume of buffer A supplemented to 0.1% (v/v) Triton X-100 detergent and incubated on ice for 10 min. The nuclei were pelleted by centrifugation (1300 x g, for 5 min, at 4°C) and washed twice with ice-cold buffer A, followed by centrifugation (1300 x *g*, 5 min at 4°C) and carefully removed the supernatant. The nuclei were resuspended in buffer A, and the concentration of nucleic acid was measured by following the protocol previously described,^47^ and then CaCl_2_ was added to 2 mM. The nuclei suspension was preincubated at 37°C for 5 min and followed by micrococcal nuclease (Worthington, cat# LS004798) digestion was carried out by following the manufacturer’s protocol. To facilitate the release of digested nucleosomes, NaCl was added to final concentrations of 200 mM, and reactions spun down at 1300 x g for 5 min at 4°C. The supernatant containing soluble nucleosomes were collected and stored at -80°C until use.

### Ni-NTA Pull-down Assays and Western Blotting

Pull-down experiments were performed by following the method previously described.^36^ 15-20 μg of acid extracted histones or nucleosomes isolated from HeLa cells were mixed with 50 μM of recombinant His_6_-tagged-BRPF2 bromodomain incubated at 4°C on a nutator for 60 min in the presence or absence of 10 μM BAY- 299 (abcam cat# ab230368) inhibitor. Samples were transferred to a fresh tube containing 50 μl of Ni-NTA agarose (QIAGEN) and incubated for 60 min at 4°C on a nutator. The beads were then washed five times with 1 ml of wash buffer containing 50 mM HEPES pH 7.5, 500 mM KCl, 2 mM EDTA, 0.1% NP-40. Bound histones or nucleosomes were eluted by incubating the beads in 20 μl of elution buffer (50 mM HEPES pH 7.5 and 500 mM Imidazole) at 4°C on a nutator. Equal volumes of eluted samples were separated on a 12%-SDS-PAGE gel and transferred onto a 0.45 μm PVDF membrane at a constant voltage of 80V for an hour at 4°C. The membrane was rinsed in TBST buffer (50 mM Tris pH 7.4, 200 mM NaCl, and 0.1% Tween-20) and blocked for an hour at room temperature (RT) in 5% milk buffer prepared in TBST. Immunoblotting was performed with the following primary antibodies: H4K5ac (Invitrogen cat# MA532009), H4K8ac (Invitrogen cat# MA533386), H4K12ac (Invitrogen cat# MA533388), and H4K16ac (Invitrogen cat# MA527794) overnight at 4°C. The membranes were washed with TBST buffer thrice at RT for five minutes each. The blots were then incubated with the HRP conjugated secondary antibodies Goat anti-Rabbit IgG (Invitrogen cat# 31466) or Goat anti-mouse IgG (Invitrogen cat# 31431) with 5% nonfat dry milk, dilution 1:20000 in TBST. The membranes were rewashed with TBST buffer thrice at RT for five minutes each. Protein bands were visualized by chemiluminescence using SuperSignal West Femto substrate (Invitrogen cat# 34094) following the manufacturer’s protocol.

### Circular Dichroism Spectroscopy

Circular Dichroism (CD) experiment was done by following the method described previously.^39^ Briefly, CD spectra were recorded on a JASCO J-1500 CD Spectrometer (JASCO, Japan) at 20°C using a quartz cell with a path length of 10 mm. Two scans were accumulated at a scan speed of 100 nm min^-1^, with data being collected at every nm from 195 to 260 nm. The BRPF2 wild-type and its mutant proteins were diluted to 2-5 μM concentartion in CD buffer containing 150 mM NaCl and 50 mM NaH_2_PO_4_ at pH 7.0. The ellipticity data was converted into molar ellipticity using the below equation.

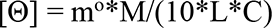

Where [Θ] is the molar ellipticity, m° is ellipticity of the sample measured, M is average molecular weight (g/mol), L is path length of the cell (cm), and C is a concentration in g/L. The protein secondary structure determination was done using K2D3 structure prediction software.^48^

### Thermal Shift Assay

The thermal shift assay is a rapid and inexpensive biochemical method often used to determine the thermal stability of protein in different *in vitro* conditions by monitoring the unfolding of the protein at increasing temperatures. The SYPRO orange fluorescent dye (Invitrogen, cat# S6650) was used to calculate the melting temperature (*T*_m_) of the wild-type BRPF2 bromodomain and its mutant proteins. The thermal shift assay was performed by following methods previously described,^39^ in a 96-well clear low-profile plate (Bio-Rad, #MLL9601) using CFX96 touch Real-Time PCR detection system. A total 25 μL assay containing 10 μg BRPF2 and its mutants were premixed with 5x SYPRO orange dye in 50 mM HEPES, pH7.5, 200 mM NaCl, and 5% glycerol. Wells containing only buffer with 5x dye were used for baseline correction. The plate was sealed with an optically clear adhesive film (Bio-Rad, #MSB1001) to prevent sample loss during heating. The temperature was gradually ramped up from 25 to 95 °C while monitoring the change in fluorescence intensity of the dye. GraphPad Prism software was used to analyze the data and calculate the melting point of the protein sample by applying nonlinear regression using the melting Boltzmann equation:

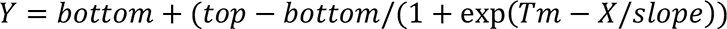

### Public ChIP-Seq Data Sets

BRPF2, BRPF3, HBO1, and input DNA ChIP-seq data in RKO cell were obtained from Gene Expression Omnibus (GEO) accession numbers: GSM1146449, GSM1556187, GSM818282, and GSM818283 respectively.^20, 23, 49^ Similarly, histone ChIP-seq raw data files of H4K5ac (GSM469975) and H4K8ac (GSM521923) in IMR90 cell and H4K12ac (GSM908963) and H3K14ac (GSM908946) in H1 cells were obtained from Gene Expression Omnibus (GEO) database.^50^

### Preprocessing and Analysis of ChIP-Seq Data

Chip seq data is analyzed using the usegalaxy server. Raw reads were quality filtered and adapter-trimmed using Trimmomatic trimming tool. Next, the trimmed reads were subsequently aligned in single-end mode to the human reference genome (hg19) using Bowtie (v. 2.2.9).^51^ Alingment followed by peak calling, is done by MACS2 (v. 2.1.0) over input.^52^ As recommended for IDR (Irreproducibility Discovery Rate) analysis, the final list of peaks were selected with an IDR threshold of 0.001 in order to obtain a conservative set of peaks. Peaks overlapping blacklisted regions as defined by the ENCODE project were discarded. All further analysis was performed in the R (v.3.4.0)/Bioconductor environment with plots generated relying on the ChIPseeker library.^53^ We used the Bioconductor package ChIPSeeker^54^ to display the overlaps between the different data sets and IGV for ChIP-seq signal visualization in genomic context. The ChIP-seq data presented in the form of line plots and heatmaps using EaSeq.^55^

## RESULTS

### The BRPF2 Bromodomain Recognizes Multiple Acetyllysine Marks on N-terminal Histone Tails

The bromodomains of BRPF2 and BRPF3 share high sequence similarity and function as part of the HBO1 HAT complex, but there is limited information on which histone ligands they bind. We recently identified multiple acetylated histone ligands recognized by the BRPF3 bromodomain using high-throughput virtual screening and *in-vitro* methods.^36^ Considering the high sequence similarity between the bromodomains of BRPF2 and BRPF3, they are likely to recognize a similar subset of acetylated histone modifications. To investigate the function of the BRPF2 bromodomain, we recombinantly overexpressed the protein in *E.coli* cells and purified it with high homogeneity (Figure S1). We performed ITC binding assays using various combinations of mono-, di- and multiple-acetylated synthesized peptides covering histone H4, H3, and H2A (Figure 1D). We found that the BRPF2 bromodomain interacts strongly with the histone H4K5ac (1-15), H4K8ac (1-15), and H4K5acK12ac (1-15), moderately with H4K12ac (1-15), H4K16ac (11-21), H4K5acK8ac (1-15), H4Kac4 (H4K5/8/12/16) (1-20), and H3K14ac (9-19) ligands, but weakly with H4K8acK12ac (1-15), and H2AK5ac (1-12) ligands (Figure 2, Figure 3 and Table 1).

**Figure 2.**
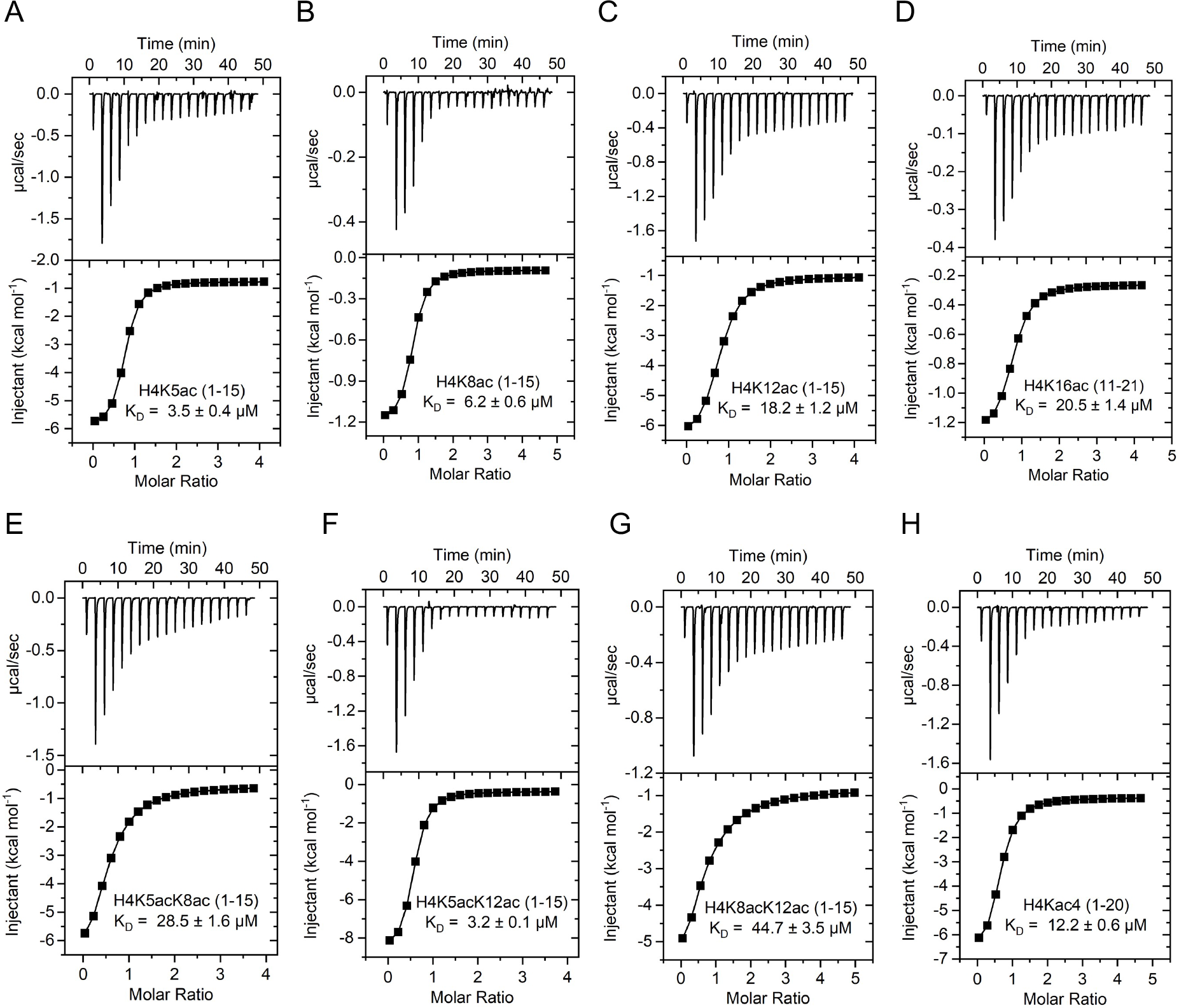
Exothermic ITC plots for binding of the BRPF2 bromodomain with different acetylated histone peptides. (A) H4K5ac (1-15), (B) H4K8ac (1–15), (C) H4K12ac (1-15), (D) H4K16ac (11–21), (E) H4K5acK8ac (1-15), (F) H4K5acK12ac (1-15), (G) H4K8acK12ac (1-15), (H) H4Kac4 (H4K5/8/12/16) (1–20). The calculated binding constants are indicated.

**Figure 3.**
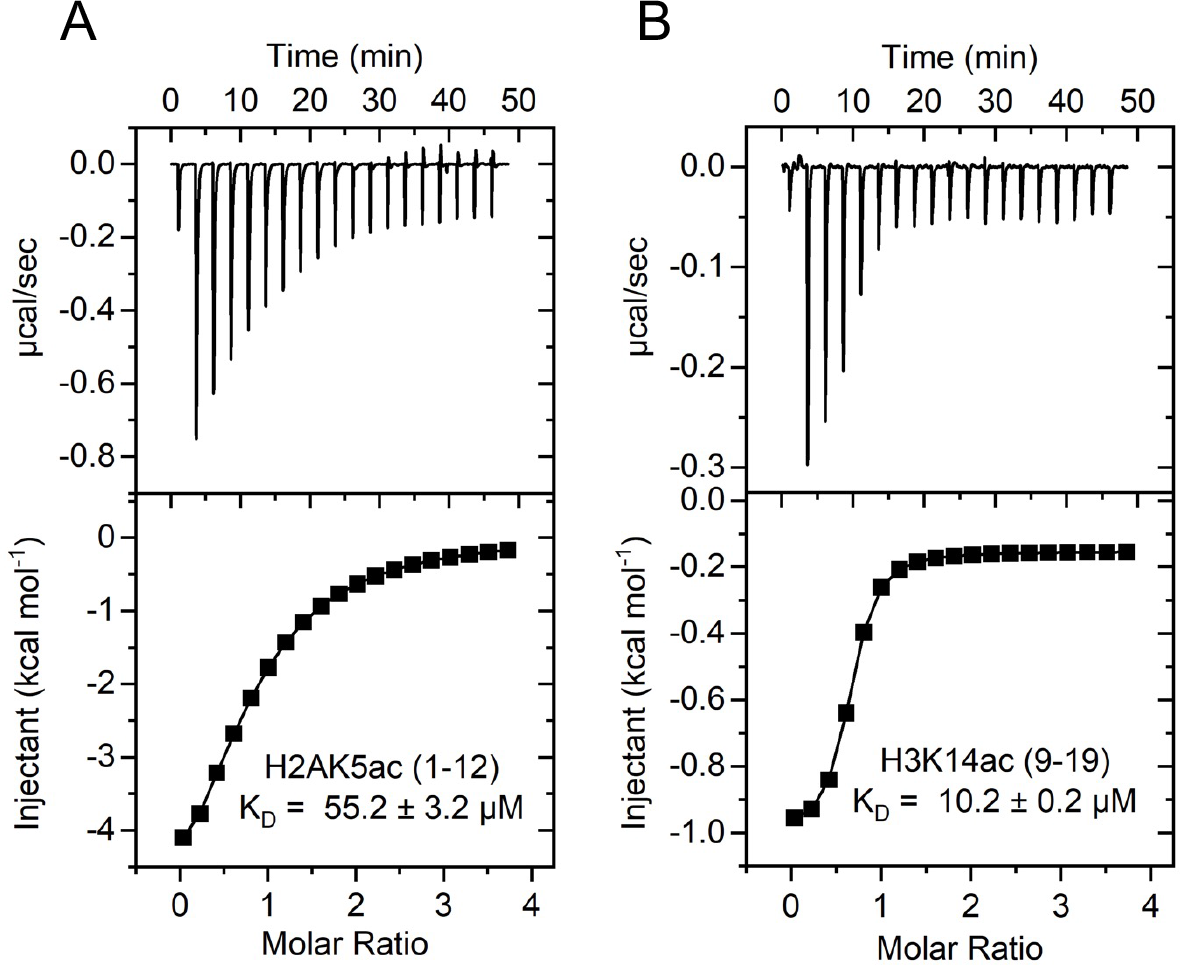
Exothermic ITC plots for binding of the BRPF2 bromodomain with histone H2A and H3 peptides. (A) H2AK5ac (1-12) and (B) H3K14ac (9–19). The calculated binding constants are indicated.

Our ITC binding results demonstrate that the BRPF2 bromodomain preferentially binds to the histone H4K5ac (1-15), H4K8ac (1-15), and H4K5acK12ac (1-15) peptides with the strongest affinities (*K*_D_ = 3.5 ± 0.4 μM, 6.2 ± 0.6 μM, and 3.2 ± 0.1 μM) respectively (Figure 2 and Table 1). The combination of H4K5acK8ac (1-15) marks decreased the BRPF2 bromodomain binding affinity about 8-fold over that of the mono-acetylated H4K5ac (1-15) mark (*K*_D_ = 28.5 ± 1.6 μM versus 3.5 ± 0.4 μM) (Figure 2 and Table 1). Notably, the combination of H4K5acK12ac (1-15) marks increased the BRPF2 bromodomain binding affinity more than 5-fold as compared to the mono-acetylated H4K12ac (1-15) ligand (*K*_D_ = 3.2 ± 0.1 μM versus 18.2 ± 1.2 μM), and also has a similar affinity compared to the H4K5ac (1-15) modification (Figure 2 and Table 1). Furthermore, other combinations of diacetylation marks H4K8acK12ac (1-15) resulted in lower binding affinities about 7-fold over H4K8ac (1-15) ligand (*K*_D_ = 44.7 ± 3.5 μM versus 6.2 ± 0.6 μM) and around 2.5-fold than the H4K12ac (1-15) ligand (*K*_D_ = 18.2 ± 1.2 μM) (Figure 2 and Table 1). Moderate affinity was observed for the mono-acetylated H4K16ac (11-21) ligand with a *K*_D_ = 20.5 ± 1.4 μM (Figure 2 and Table 1). Next, we tested tetra-acetylated H4Kac4 peptide carrying the acetylation sites at K5/8/12/16 on N-terminal tails of histone H4. We found that the BRPF2 bromodomain bound to H4Kac4 (1-20) peptide with a dissociation constant of *K*_D_ = 12.2 ± 0.6 μM (Figure 2 and Table 1). These results suggest that the BRPF2 bromodomain recognizes multiple acetylation marks on histone H4, with the singly acetylated mark K5ac and K8ac recognition driving the binding interaction. While the combination modification like diacetylated H4K5acK8ac or H4K8acK12ac that are separated by 3-4 residues apart is inhibitory, possibly due to steric hindrance during binding with the bromodomain. And more importantly the diacetylation mark H4K5acK12ac which is separated by 7 residues showed similar binding to that of H4K5ac mark again confirming that singly acetylated K5 is driving the binding interaction.

More recent studies on the family IV bromodomains revealed that BRPF1 recognizes H3K14ac and H2AK5ac ligands with a moderate affinity, and the BRPF3 and ATAD2/B forms transient interactions with the histone H2AK5ac ligand.^34–36, 56, 57^ We also tested the binding affinity of the BRPF2 bromodomain with histone peptides H3K14ac (9-19) and H2AK5ac (1-12). We found that the BRPF2 bromodomain binds to H3K14ac and H2AK5ac peptides with dissociation constants (*K*_D_ = 10.2 ± 0.2 μM and 55.2 ± 3.2 μM) respectively (Figure 3, Table 1). No binding was observed in the case of unmodified histone H4 (1–20), H3 (1–24), and H2A (1– 12) peptides (Figure S2). Collectively, our ITC binding results demonstrate that the BRPF2 bromodomain strongly recognizes multiple acetyllysine residues on the N-terminal tails of histone H4.

### MD Simulations Predict the Acetyllysine Binding Pocket Residues

We carried out molecular dynamics (MD) simulations to understand the mode of acetyllysine binding and stability of the bromodomain-histone peptide complexes. Fitst, we covalently modified the selected lysine residues on the N-terminal tails of histone H4 using an in-silico approach. Next, we carried out flexible docking studies for the BRPF2 bromodomain with unacetylated and acetylated histone H4 (H4K5ac and H4K5acK12ac) ligands. The best ranked binding conformations of histone ligands obtained from docking studies were used to perform the MD simulations. The RMSD (Root mean square deviation) was used to analyze the stability of the bromodomain and histone peptide complexes. RMSD results revealed that both acetylated ligands H4K5ac and H4K5acK12ac are stable throughout the simulations, whereas unacetylated H4 ligand has significantly deviated within 100 ns of MD simulation (Figure 4A). Similarly, the analysis of RMSF (Root mean square fluctuation) for each residue demonstrates that unacetylated H4 ligand showed significant fluctuations in the ZA loop of the bromodomain compared to the acetylated H4 ligands (Figure 4B). These results highlight the importance of the ZA loop in acetyllysine recognition and stabilizing the interactions between the bromodomain and acetylated histone peptide complexes.

**Figure 4.**
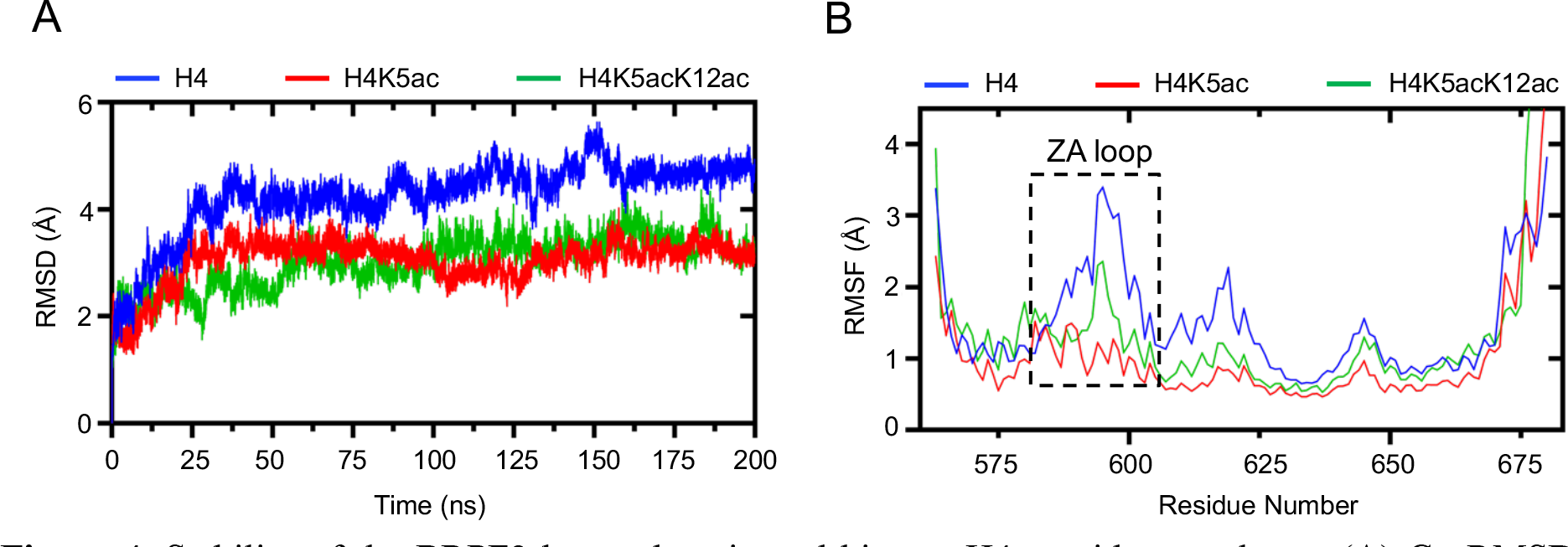
Stability of the BRPF2 bromodomain and histone H4 peptide complexes. (A) Cα RMSD demonstrates the stability of the protein backbone during the MD simulation. (B) The Cα carbon RMSF of the BRPF2 bromodomain is plotted against the residue numbers. The BRPF2-histone H4K5ac and H4K5acK12ac peptide complexes are showed very similar fluctuations, but the BRPF2-unacetylated H4 peptide display observable changes in the ZA-loop region.

Next, the visual analysis of MD trajectories revealed that the K5ac residue in two of the histone peptides, H4K5ac and H4K5acK12ac inserts deep into the binding cavity of the BRPF2 bromodomain (Figure 5A,B). Further, the analysis of MD trajectories using the PLIP (Protein-Ligand Interaction Profiler) server demonstrates that coordination of the K5ac residue of H4K5ac and H4K5acK12ac peptides occurs through a combination of hydrogen bonds and hydrophobic interactions with the bromodomain binding pocket residues (Figure 5A,B and Table S3 and S4). Specifically, the K5ac residue of the H4K5ac peptide forms a hydrogen bond interaction with the critical acetyllysine binding pocket residue N642 and indirectly interacts with the Y599 residue of bromodomain through water bridges (Figure 5A and Table S3). Similarly, the K5ac residue of the H4K5acK12ac peptide interacts with two critical binding pocket residues, N642 and Y599, of the BRPF2 bromodomain through hydrogen bonding and water coordination, respectively (Figure 5B and Table S4). In addition, the K5ac residue of the H4 peptides makes hydrophobic interactions with residues F587 and F648 of the BC loop region of the bromodomain. Together, these results correspond to the ITC binding studies conducted with the multiple acetylated histone H4 peptides, and the K5ac recognition mainly drives the binding interaction with the BRPF2 bromodomain.

**Figure 5.**
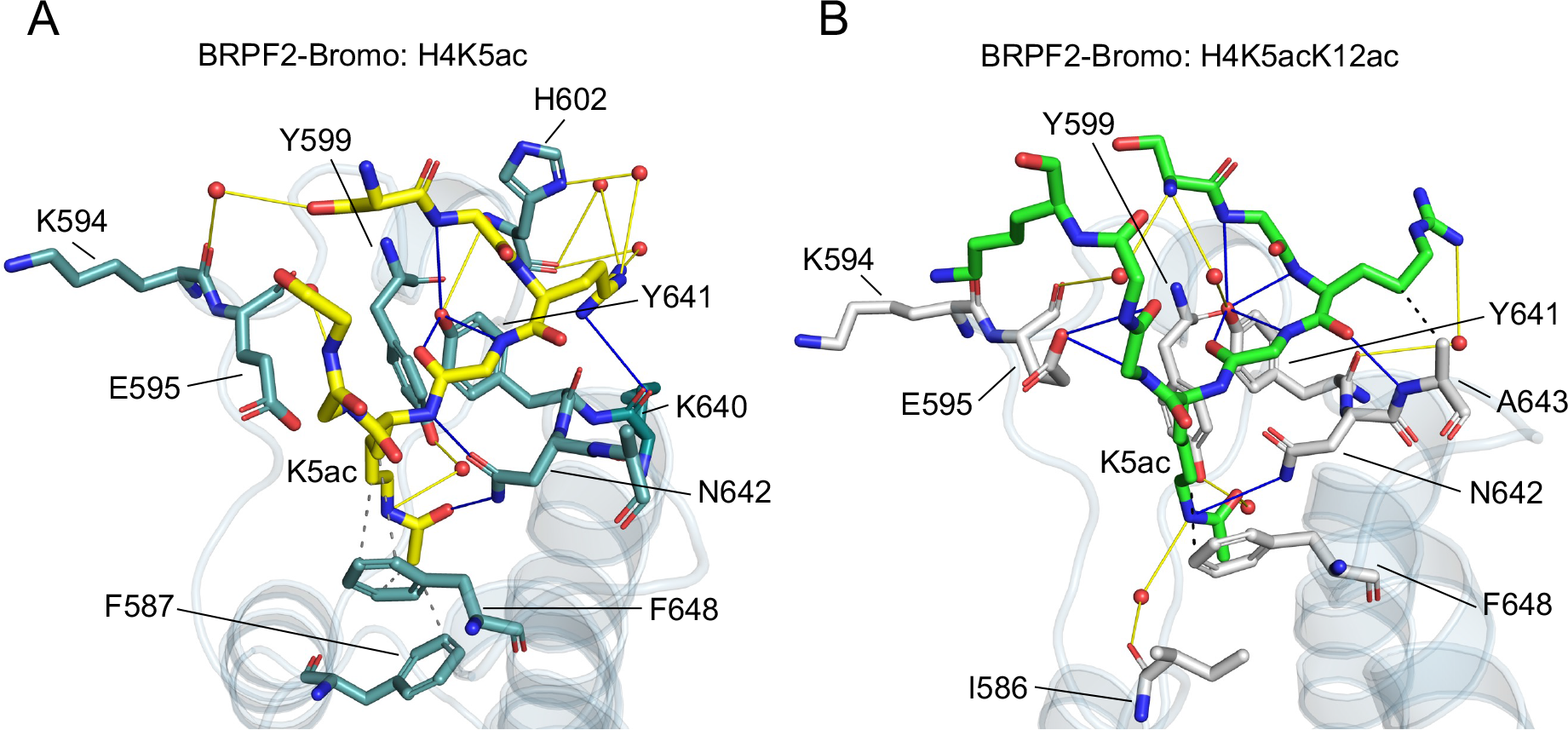
MD models showing the dynamics of acetyllysine binding in the bromodomain binding pocket. (A,B) Cartoon representation shows the molecular contacts between the BRPF2 bromodomain and acetylated histone peptides H4K5ac and H4K5acK12ac. The dotted and solid lines represent the hydrogen bonding and hydrophobic interactions, respectively.

### BRPF2 Bromodomain Recognizes Endogenous Acetylated Histone H4

To address whether the BRPF2 bromodomain binds to endogenous acetylated histone H4, we performed pull-down assays by incubating the recombinantly produced 6xHis-tagged BRPF2 bromodomain with hyperacetylated H4 isolated from human cells (Figure 6A and Figure S3). Subsequent enrichment with Ni-NTA beads and probing of the bound material by western blot against anti-H4ac antibodies revealed that the BRPF2 bromodomain efficiently binds to endogenous histone H4K5ac and H4K8ac as compared to H4K12ac and H4K16ac marks (Figure 6B and Figure S4A). Notably, the BRPF2 inhibitor BAY-299 reduced the interaction approximately 3-fold between the BRPF2 bromodomain and acetylated histone H4, as probed with anti-H4K5ac antibody (Figure 6C and Figure S4B). We carried out computational molecular docking studies to further investigate the mode of inhibitor binding in the bromodomain binding pocket. Our docking results revealed that BAY-299 preferably occupies the acetyllysine binding pocket of the BRPF2 bromodomain and forms a hydrogen bond with the anchor (N642) residue (Figure 6D), which is a highly conserved residue throughout the bromodomain family and critical for acetyllysine recognition.^10^ Unfortunately, no antibodies are currently available that specifically recognize diacetyllysine marks on histone H4. Together, these results demonstrate that the BRPF2 bromodomain can recognize the identified histone H4 marks at the endogenous histone level.

**Figure 6.**
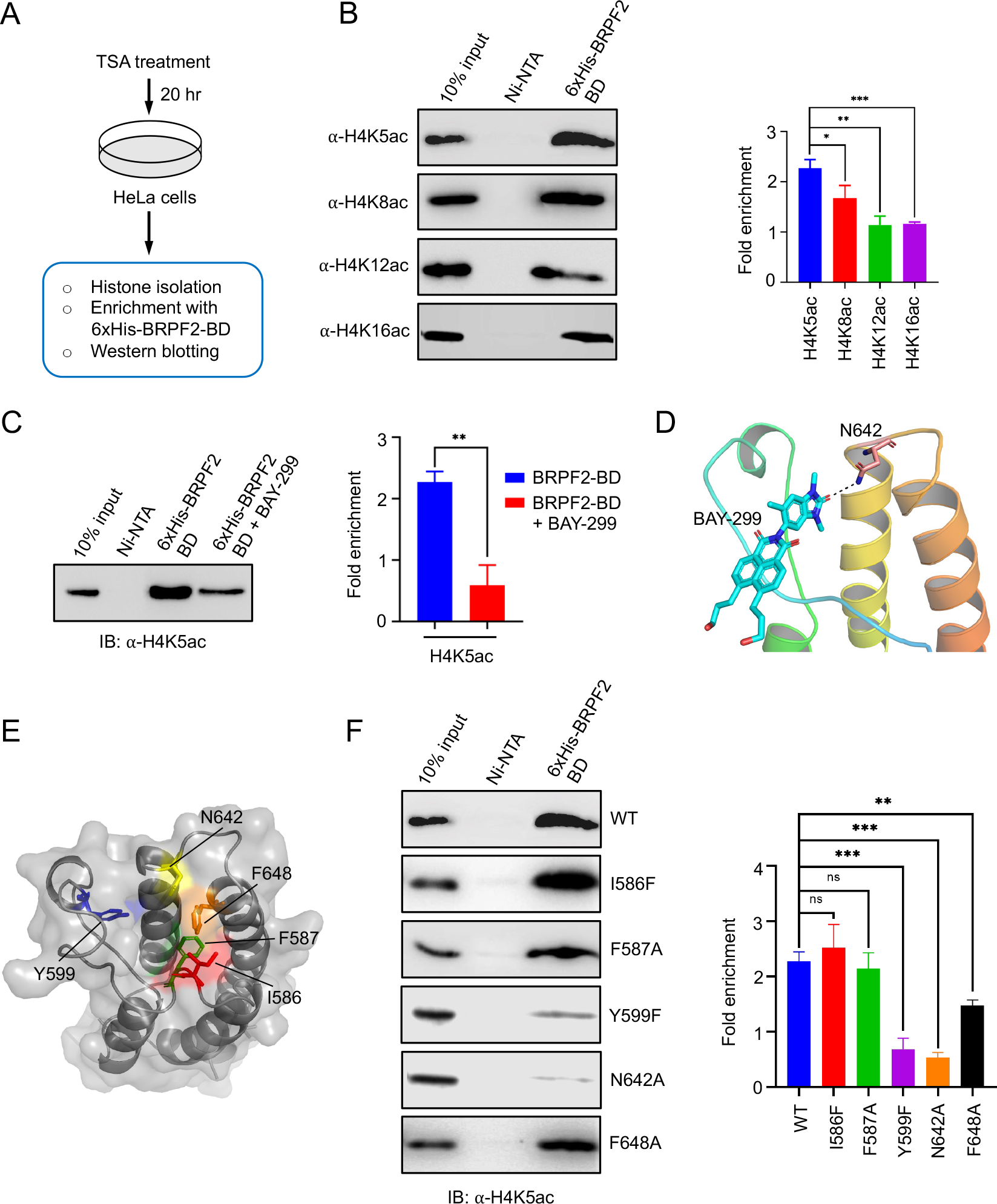
Interaction of BRPF2 bromodomain wild-type and its mutants with endogenous histone H4 (A) Hyperacetylated histone H4 was purified from HeLa cells. (B) The BRPF2 bromodomain was incubated with hyperacetylated H4 and bound acetylated H4 was enriched by Ni-NTA bead, followed by western blot against anti-acetylated H4 antibodies. (C) The BRPF2 bromodomain was incubated with hyperacetylated H4 in the presence or absence of BAY-299 inhibitor, and bound acetylated H4 was enriched by Ni-NTA bead, followed by western blot against anti-H4K5ac antibody. (D) The cartoon representation shows that the docked conformation of BAY-299 forms a hydrogen bond with N642 residue of the BRPF2 bromodomain. (E) The transparent cartoon-surface representation of the BRPF2 bromodomain shows the highlighted amino acid residues are subjected to mutagenesis. (F) The mutational analysis of acetyllysine binding pocket residues demonstrates that the Y599F and N642A mutants significantly abolished the interaction of BRPF2 bromodomain with H4K5ac modification as compared to the other mutants. The relative band intensities were measured by ImageJ and normalized for each immunoblotting. Bars represent mean ± SD derived from n=3 individual experiments. Single star indicates p-value ≤ 0.05, double star indicates p-value ≤ 0.01, and triple star indicates p-value ≤ 0.001.

### Mutational Analysis Identifies Bromodomain Binding Pocket Residues that are Critical for Acetyllysine Recognition

Next, we sought to investigate the amino acid residues important for bromodomain-acetyllysine recognition. With the help of our MD simulation studies, we specifically targeted five amino acid residues for the mutational analysis (Figure 6E). We chose three of the five residues from the ZA loop region, namely, the hydrophobic amino acid residues I586 and F587, which are part of the RIF shelf similar to the WPF shelf in BET bromodomains, and Y599, which faces the acetyllysine binding pocket of the BRPF2 bromodomain. The remaining two residues, N642 and F648, were selected from the BC loop region. We mutated these residues to alanine, except two residues, I586 and Y599, which were mutated to phenylalanine. Using a bacterial expression system, we expressed all the five mutants and purified them with high homogeneity for the binding studies (Figure S5). We carried out pull-down assays by incubating the BRPF2 bromodomain mutants with the endogenous hyperacetylated histone H4. The bound proteins were enriched with Ni-NTA beads, followed by western blot with anti-H4K5ac antibody revealed that the Y599F and N642A mutants significantly abolishes the interaction with the histone H4K5ac mark (Figure 6F and Figure S6). We also observed that the F648A mutation reduced the binding approximately 1.5-fold, and F587A showed a minimal binding effect, whereas I586F enriched the H4K5ac mark similar to the wild-type protein (Figure 6F and Figure S6). These results closely support the ITC binding studies carried out for the BRPF2 bromodomain mutants I586F, F587A, Y599F, N642A, and F648A with the histone H4K5ac (1-15) peptide (Figure 7 and Table 2).

**Figure 7.**
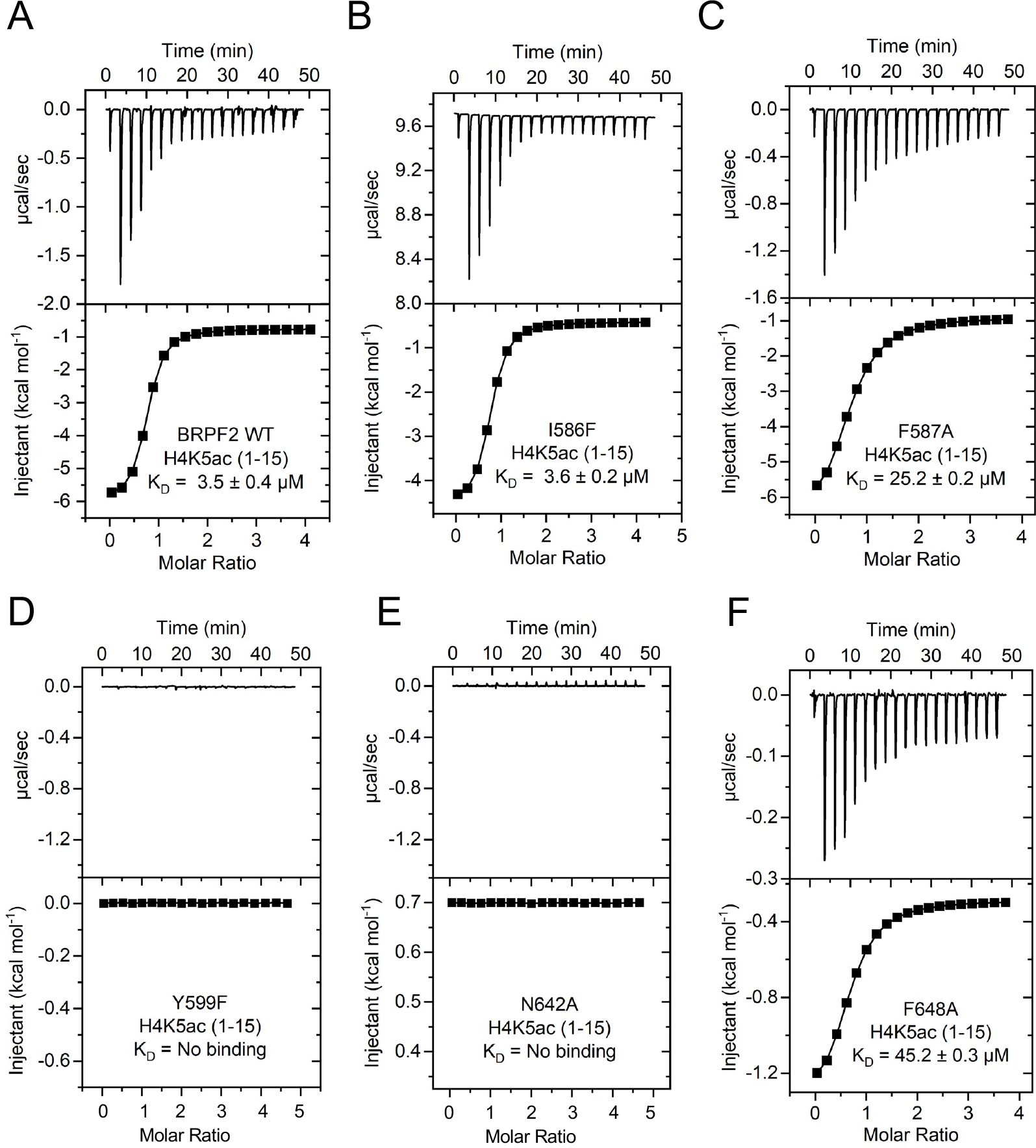
Exothermic ITC plots for binding of the wild-type BRPF2 bromodomain and its mutants with histone H4K5ac (1-15) peptide. The calculated binding constants are indicated.

Next, we performed circular dichroism (CD) experiments for the BRPF2 bromodomain mutants to verify whether the loss of binding is due to any conformational changes in the protein after the mutation. Analysis of the BRPF2 bromodomain wild-type and its mutants revealed no significant alterations in the bromodomain secondary structure (Figure 8A and Table S5). We further characterized the melting temperature (*T*_m_) of the BRPF2 bromodomain mutants by a thermal shift assay. We found that the Y599F and N642A mutants displayed significantly lower *T*_m_ than the wild-type protein (Figure 8B), suggesting that these mutants are less stable than the wild-type protein. In contrast, the F648A mutant showed a higher stabilization profile.

**Figure 8.**
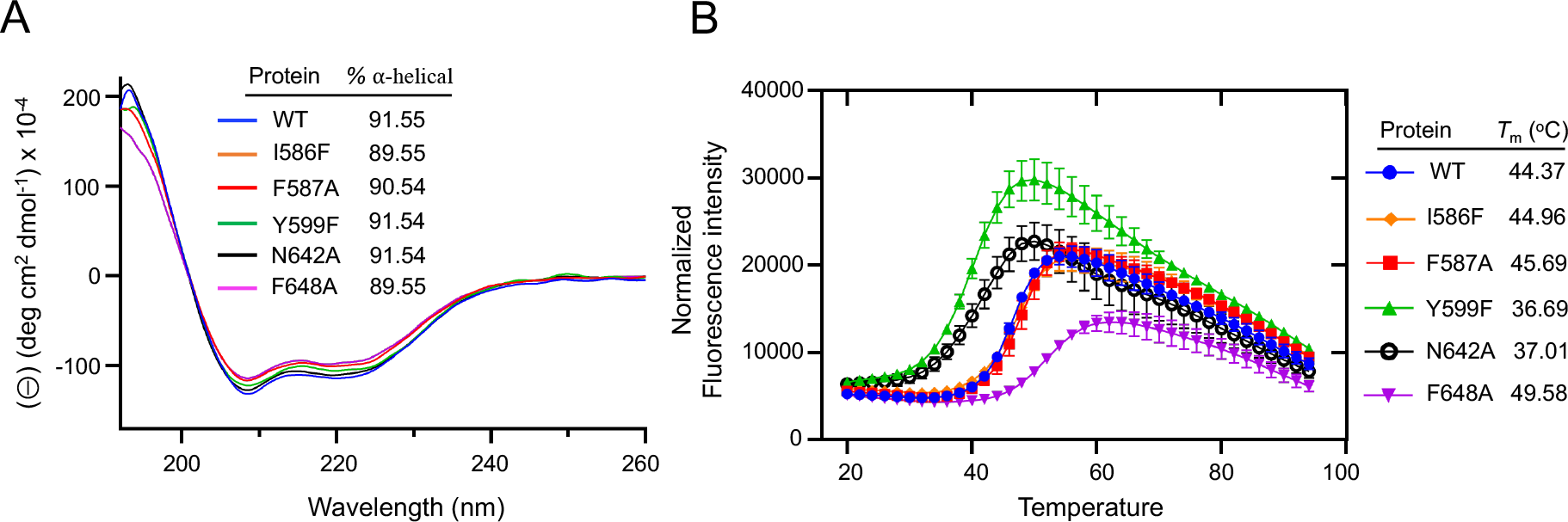
Circular dichroism spectra and thermal stability assay (TSA) of the wild-type and mutant BRPF2 bromodomains. (A) Secondary structure of the BRPF2 bromodomain wild-type and its mutants measured by circular dichroism. The percentage of ⍺-helical content for each bromodomain is listed in the inset. (B) Thermal stability assay shows the unfolding curve for the BRPF2 bromodomain wild-type and its mutant proteins.

Conversely, the other two mutants, I586F and F587A, showed similar *T*_m_ values as compared to the wild-type. Overall, these *T*_m_ values are in good agreement with the binding affinities of the mutants for H4K5ac marks in the ITC and pull-down experiments (Figure 6F, and Figure S6). Collectively, these results suggest that the ZA loop and BC loop of the BRPF2 bromodomain plays a vital role in stabilizing the overall fold of the bromodomain and acetyllysine recognition.

### BRPF2 Bromodomain Strongly Recognizes H4K5ac marks at the Mononucleosome Level

To address whether the BRPF2 bromodomain prefers to recognize identified histone H4 marks bearing the nucleosomes, we isolated mononucleosomes using MNase fragmented chromatin derived from HeLa nuclei (Figure 9A and Figure S7). We then performed pull-down experiments on the mononucleosomal pools employing 6xHis-tagged BRPF2 bromodomain and subsequently enriched them with Ni-NTA beads, followed by western blot with anti-acetylated H4 antibodies. Notably, we found that the H4K5ac mark associates more robustly with the BRPF2 bromodomain as compared to H4K8ac, H4K12ac, and H4K16ac marks at the mononucleosomal level (Figure 9B and Figure S8). Furthermore, we also observed substantial enrichment of the H4K8ac marks by the BRPF2 bromodomain, and these observations are consistently supplemented by the pull-down assays conducted with the endogenous acetylated histone H4 (Figure 6B). We interpret the enhanced binding of H4K5ac and H4K8ac bearing-mononucleosomes to suggest that both marks may play a role in nucleosome-level binding, yet the H4K5ac interaction is dominant. However, future experiments with semi-synthetic nucleosomes with defined acetylation modifications will help resolve the inter versus intra nucleosomal binding.^45^ We observed minimal binding of H4K12ac and H4K16ac marks to the BRPF2 bromodomain. Together, these results are closely supported by our ITC binding and pull-down assays carried out for the BRPF2 bromodomain with H4 peptides and endogenous histone H4, respectively (Figure 2 and Figure 6B).

**Figure 9.**
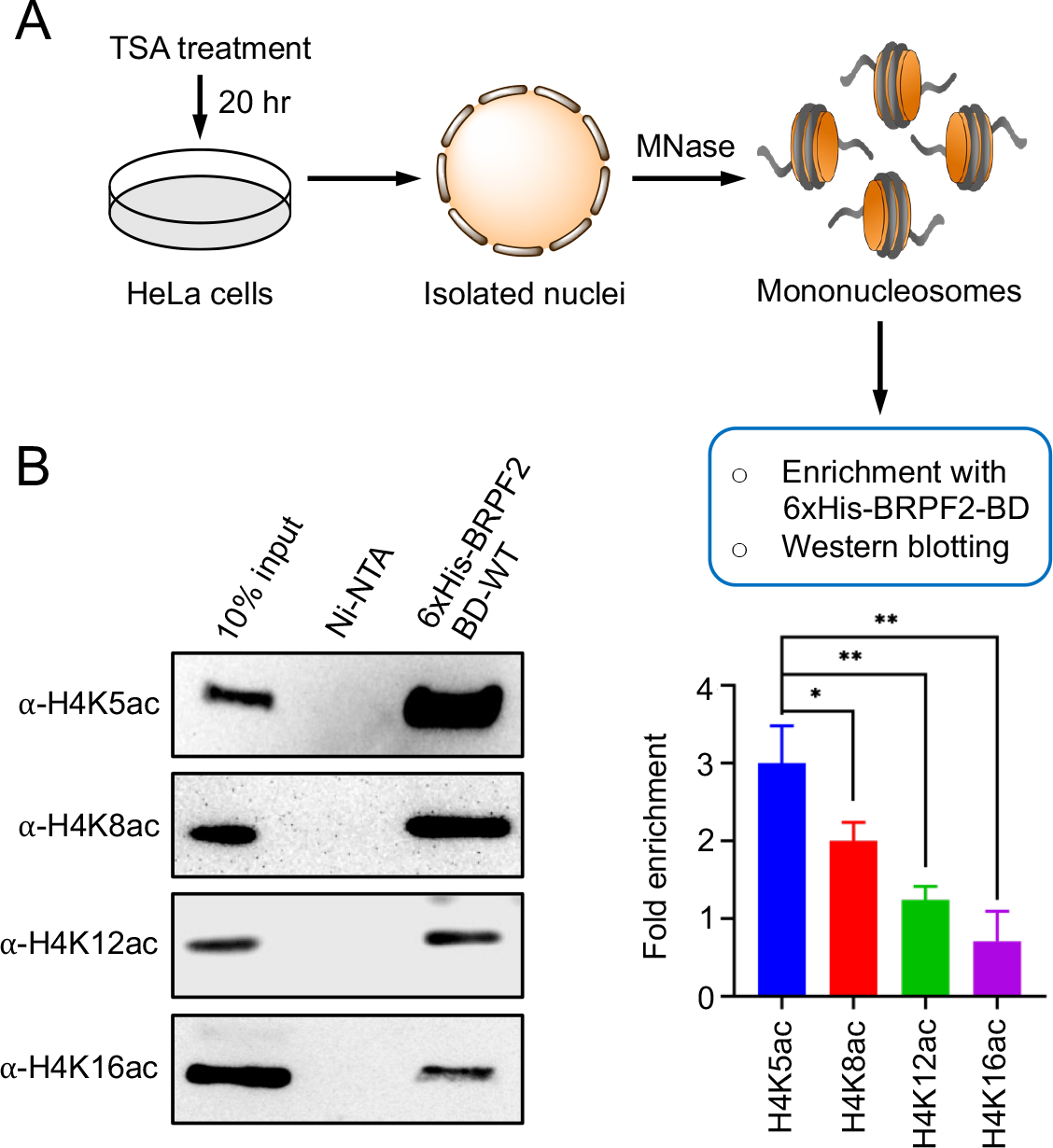
The BRPF2 bromodomain strongly recognizes the mono-acetylated H4K5ac mark at the mononucleosome level. (A) Schematic representation showing the isolation of mononucleosomes from HeLa cells. (B) Recombinantly produced 6xHis-tagged BRPF2 bromodomain was used to pull-down native mononucleosomes, and the bound material was probed by western blot against the anti-acetylated H4 antibodies. The relative band intensities were measured by ImageJ and normalized for each immunoblotting. Bars represent mean ± SD derived from n=3 individual experiments. Single star indicates p-value ≤ 0.05 and double star indicates p-value ≤ 0.01.

### BRPF2 Associates with the Genome Extensively

To analyze genome-wide occupancy of BRPF2, we analyzed published chromatin immunoprecipitation sequencing (ChIP-seq) data from Gene Expression Omnibus (GEO) in the RKO cell line. Raw data were trimmed and mapped onto the Human Genome version 19 (hg19) as a reference genome. ChIP-seq analysis revealed extensive occupancy of BRPF2 on the genome (Figure 10A). Over 18900 recovered BRPF2 peaks were consistent across the Irreproducible Discovery Rate (IDR) analysis. We observed BRPF2 occupancy near the annotated transcription start sites (TSS), with over 80% peaks occurring at upstream promoter regions (Figure 10B). Examining the average BRPF2 occupancy profile near all TSS, we found a conserved pattern featuring bimodal occupancy of BRPF2 with peaks flanking the TSS, with a more substantial peak typically occurring downstream of the TSS (Figure 10C). There is enrichment near the TSS and not at the transcription termination site (TTS) along the entire transcribed loci of gene bodies (Figure 10D).

**Figure 10.**
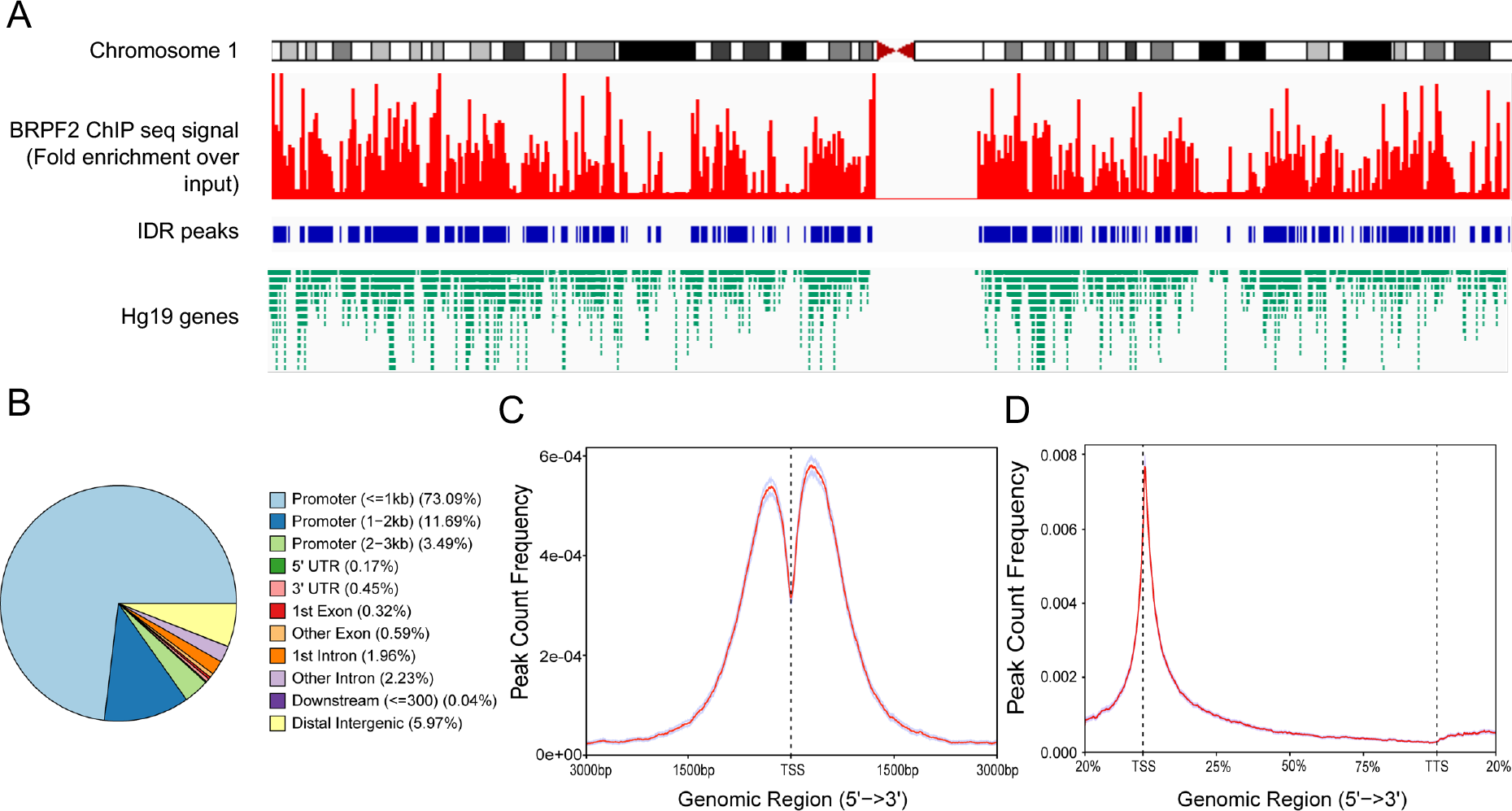
ChIP-seq analysis shows extensive occupancy of BRPF2 along the genome. (A) ChIP-seq signal (fold enrichment over input) of BRPF2 shown along the chromosome 1 along with peaks determined using the irreproducibility discovery rate (IDR). Known Hg19 genes and transcripts are aligned. (B) Pie chart representation of BRPF2 binding sites in the genome. (C) Average profile of ChIP peaks of BRPF2 binding to transcription start site (TSS) region. (D) ChIP peaks of BRPF2 binding to the gene bodies from the transcription start site (TSS) to the transcription termination site (TTS).

### BRPF2 Co-localizes with the H4K5ac and H4K8ac Marks in the Genome

Based on our biochemical experiments, we envisioned that BRPF2 localizes along with HBO1 at the different histone H4 marks in the genome. In our previous study, we showed that the BRPF3 bromodomain also recognizes similar set of histone marks as that of BRPF2.^36^ To gain molecular insight into the differential localization with these histone tags of the BRPF paralogues, we included BRPF3 in this study. To investigate this, we performed ChIP-seq data analysis from published data sets of BRPF2 (GSM1146449), BRPF3 (GSM1556187), and HBO1 (GSM818282) and the three most crucial histone H4 marks that strongly binds to the BRPF2 bromodomain viz., H4K5ac (GSM469975), H4K8ac (GSM521923) and H4K12ac (GSM908963) as well as H3K14ac (GSM908946) as a reference mark which is reported to colocalize with HBO1 and BRPF3.^23^ After carefully analyzing the data, we found a strong correlation between the occupancy of BRPF2, BRPF3, HBO1 and histone marks (Figure 11). We found different regions on the genome where BRPF2, BRPF3, HBO1 and H4K5ac, H4K8ac, H4K12ac and H3K14ac colocalizes in the transcription start site of the following genes namely *HOXA4*, *CBX3*, *HNRNPA2B*, *AIFM2* and *TYSND1* (Figure 11A). Next, we performed Venn diagram analysis to understand the overlap of the BRPF2 associated genes with histone marks H4K5ac, H4K8ac, H4K12ac and H3K14ac, and BRPF3 and HBO1. We found that 5522 (29.2%) peaks of BRPF2 overlaps with the peak of histone marks H4K5ac, H4K8ac, H4K12ac and H3K14ac as illustrated in the Venn diagram (Figure 11B). We further analysed the occupancy of H4K5ac and H4K8ac histone marks in the BRPF2, BRPF3 and HBO1 associated genes and found that they overlapped with 7320 (39.5%) of the BRPF2 peaks (Figure 11C). Moreover, additional analysis performed using EaSeq to generate the line and heat map plots further demonstrate H4K5ac, H4K8ac, H4K12ac and H3K14ac localization at the promoters of BRPF2 occupied gene regions (n=14970) (Figure 11D). From this analysis we found that H4K5ac, H4K8ac and H3K14ac marks occupancy is more abundant than the H4K12ac mark throughout the BRPF2 localized regions. Notably, we found that the BRPF3 and HBO1 also colocalize with the BRPF2 associated genes in the genome (Figure 11D).

**Figure 11.**
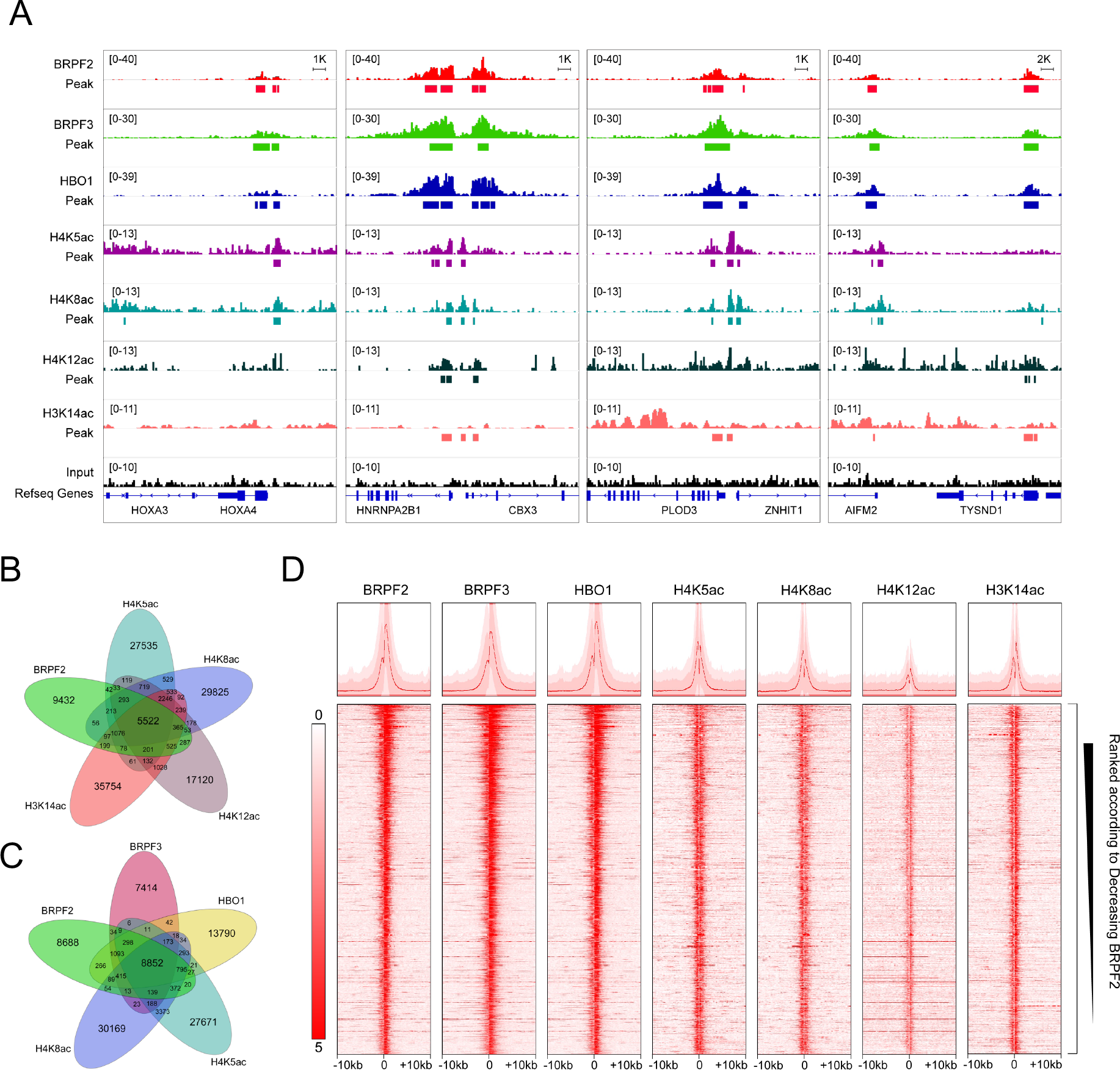
ChIP-seq analysis of BRPF2 binding to the human genome. (A) BRPF2 enrichment (over input) signal obtained by ChIP-seq (GSM1146449) analysis and compared with the signals of HBO1 (GSM818282), BRPF3 (GSM1556187) in RKO cells, H4K5ac (GSM469975), H4K8ac (GSM521923) in IMR90 cells and H4K12ac (GSM908963) and H3K14ac (GSM908946) in H1 cells. (B) Venn diagrams illustrating genomic binding sites overlap between BRPF2 and histone tags H4K5ac, H4K8ac, H4K12ac and H3K14ac. (C) Venn diagrams showing overlap of genome binding sites between BRPF2, BRPF3, HBO1 and histone marks H4K5ac, H4K8ac. (D) (Bottom) Heat maps of BRPF2, BRPF3, HBO1, H4K5ac, H4K8ac, H4K12ac and H3K14ac ChIP-seq signal to +/-10 kb around the TSS of BRPF2-localized genes. (Top) Analysis of signals around the gene transcription start sites of BRPF2 localized genes shown in line graphs observed in ChIP-seq datasets. Genes were sorted from high to low by the BRPF2 peak signal.

**Figure 12.**
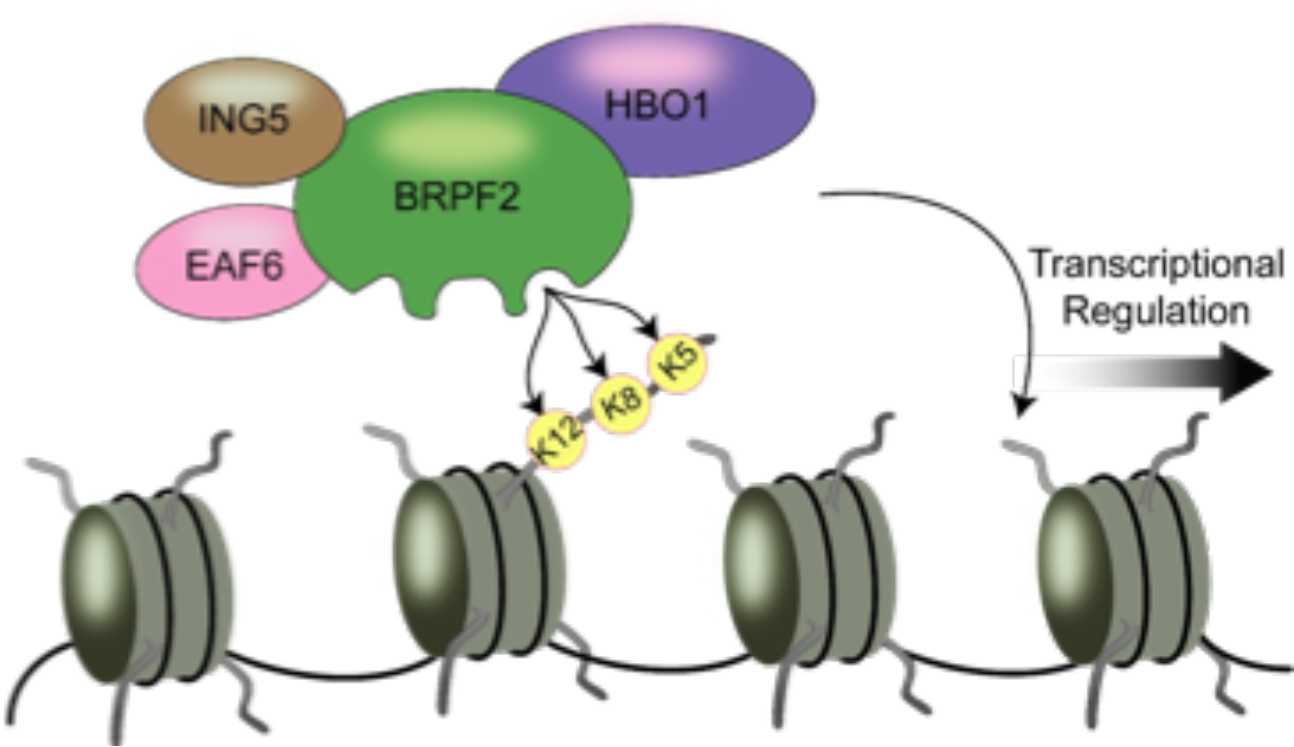
Schematic model depicting the recruitment of BRPF2-HBO1 complex to chromatin for the downstream transcriptional regulation.

## DISCUSSION

The BRPF2 and BRPF3 scaffolding proteins are important for the HAT activity of HBO1, which acetylates histone H3 and H4.^18, 19^ These scaffolding proteins regulate the HAT activity of HBO1 towards H3K14, and the deletion of HBO1 in mice exhibited a profound reduction of global H3K14 acetylation causing defects in embryonic development.^19^ Similarly, the loss of BRPF2 in mice led to a decreased histone H3 lysine 14 acetylation at the promoters of erythroid developmental regulator genes.^21^ However, it is not clear how BRPF2 directs the recruitment of HBO1 HAT to chromatin to regulate the gene transcription. Recent studies on the BRPF3 bromodomain demonstrate that it recognizes multiple acetylated lysine residues on the N-terminal tails of histone H4 and shows preferential interactions with H4K5ac and H4K5acK12ac marks.^36^ Since the sequence similarity between the BRPF2 and BRPF3 bromodomains is high, most likely, the BRPF2 bromodomain recognizes a similar set of acetylated histone modifications. As expected, we found that the BRPF2 bromodomain has a similar preference for selecting acetyllysine modifications on the histone H4 tail. Additionally, we found that the BRPF2 bromodomain strongly binds to mono acetylated histone H4 and H3 ligands (H4K8ac, H4K12ac, H4K16ac, and H3K14ac), and multiple acetylated H4 ligands (H4K5acK8ac, H4K8acK12ac, and H4Kac4) as compared to the BRPF3 bromodomain.^36^ Collectively, our ITC binding studies suggest that the BRPF2 and BRPF3 bromodomains have slightly different preferences for binding to the acetylated histone ligands due to the variability in the ZA and BC loop regions of the bromodomain.

Flexible ligand docking followed by MD simulation studies were used to model the BRPF2 bromodomain and histone H4 peptide complexes. Our MD models suggest that hydrophobic interactions between the residues F587, A643, and F648 of the bromodomain and histone peptides drive acetyllysine binding. Once inserted into the binding cavity, acetyllysine makes strong hydrogen bonds with Y599 and N642 residues. These residues are part of the ZA and BC loop regions, are highly conserved throughout the bromodomain family, and play a critical role in acetyllysine recognition.^10, 11^

Analysis of the BRPF2 bromodomain mutants coupled with pull-down and ITC binding assays identified the critical residues involved in the acetyllysine binding. Both enrichment and ITC binding studies revealed that the I586F mutation in the “RIF shelf” region of the BRPF2 bromodomain did not impair acetyllysine binding, instead it strongly binds to H4K5ac mark similar to the wild-type efficiency. Moreover, previous studies have shown that I586F mutation in the bromodomain of BRPF2 permitted the selection of larger acylation marks like butyrylation without affecting acetyllysine binding.^37^ Notably, the Y599F and N642A mutations completely abolished the binding interaction between the BRPF2 bromodomain and the H4K5ac ligand, due to the loss of water coordination and direct hydrogen bonding with acetyllysine residue in Y599F and N642A mutants, respectively. Thus, the ordered water molecules in the bromodomain binding pocket and conserved asparagine “anchor” residue is critical for recognizing acetylated histone ligands.^58, 59^ Recent studies have extensively characterized the histones targets of the family IV bromodomains, including BRPF1, BRPF3, ATAD2, and ATAD2B bromodomains. These proteins have a preference for selecting the histone H4K5ac and H4K5acK12ac ligands with varying binding affinities.^35, 36, 56, 57^ Importantly, our study shows that the BRPF2 bromodomain has a similar preference for selecting the histone H4K5ac and H4K5acK12ac ligands with a higher affinity.

Although many studies focus on individual histone PTMs in isolation, recent studies have shown that the BPTF bromodomain strongly discriminates between acetylated histone H4 marks at the peptide versus nucleosome level.^45^ We also characterized the interactions between the BRPF2 bromodomain and histone H4 marks at the nucleosome level. We found that the BRPF2 bromodomain bound a greater quantity of H4K5ac mononucleosomes as compared to the other histone H4 modifications. Overall, our nucleosomal enrichment data demonstrate that the H4K5ac mark is the preferred histone target of the BRPF2 bromodomain.

The histone binding property of bromodomains tethers them to the chromatin to recruit regulatory complexes for transcription regulation.^60^ Our analysis of published ChIP-seq data revealed that BRPF2 extensively associates with the genome and mostly occupy the promoter region showing a bimodal distribution pattern around the TSS. Previous reports have shown that similar to BRPF2, bromodomain containing proteins like BRD2, BRD3 and BRD4 also shows a bimodal signature of binding around the TSS.^61^ ChIP-seq analysis further supports our *in vitro* ligand binding data, demonstrating that BRPF2 along with HBO1 HAT localizes at different acetylated H4 regions in the genome showing a strong occupancy with the H4K5ac and H4K8ac marks followed by H4K12ac mark. As expected, BRPF2 also showed a strong co-occupancy with the H3K14ac mark which is a substrate of the HBO1 HAT.^23^ At the BRPF2 localized regions, HBO1, H4K5ac, H4K8ac, H4K12ac and H3K14ac shows a bimodal distribution pattern around the TSS similar to that exhibited by RNA Pol II, where the first peak demonstrates the instance of Pol II positioning and the second peak reflects pausing at the +1 nucleosome during transcription.^62^ Altogether, we identified novel acetylated histone ligands of BRPF2 and our study provide insights into how binding of BRPF2 with acetylated histone recruits HBO1 for transcription regulation.

## CONCLUSION

This study discovered novel histone ligands of the BRPF2 bromodomain that specifically recognizes different acetyllysine marks on the N-terminal tails of histone H4, H3, and H2A. We identified that the mono-(H4K5ac, H4K8ac) and di- (H4K5acK12ac) acetylated histone peptides are the preferred targets of the BRPF2 bromodomain. We establish that the BRPF2 bromodomain strongly binds to H4K5ac marks on histone peptides, endogenous histones, and mononucleosomal levels. Notably, it has been shown that the BRPF2-HBO1 co-localizes in the genome and shares a significant portion of their target genes involved in transcriptional regulation.^23^ Additionally, BRPF2-HBO1 interaction is critical for the proper functioning of HBO1, particularly in the global acetylation of H3K14.^23^ Thus, our findings offer a potential molecular mechanism that the BRPF2 bromodomain recruits the HBO1 HAT to chromatin by recognizing its histone interacting partners for the downstream transcriptional regulation.

## ACCESSION CODES

BRPF2 (BRD1): O95696 (UniProt ID)

## ASSOCIATED CONTENT

### Supporting Information

The Supporting Information is available free of charge on the ACS Publications website at DOI: The Supporting Information contain peptide characterization including ESI-MS peptide data, HPLC purity traces for peptides, ITC binding data, Figure S1-S34 and Table S1-S5.

## AUTHOR INFORMATION

**Corresponding Author**:*

**ORCID**: Babu Sudhamalla: 0000-0002-6610-1424.

## AUTHOR CONTRIBUTIONS

B.S. conceived the ideas. S.B., A.R., and B.S. designed the experiments. S.B., A.R., and J.P. performed the biochemical experiments. S.B. carried out the docking and MD simulations studies. S.B., AR., J.P., and B.S. analyzed data. B.S. wrote the paper.

## Notes

The authors declare no competing financial interest.

## ACKNOWLEDGMENTS

The authors thank the research funding from IISER Kolkata, infrastructural facilities supported by IISER Kolkata, and DST-FIST (SR/FST/LS-II/2017/93).

## ABBREVIATIONS

BRD: Bromodomain
ITC: Isothermal titration calorimetry
MD: Molecular dynamics
BRPF1/2/3: Bromodomain and PHD finger containing protein 1/2/3
PHD: Plant homeodomain
ATAD2B: ATPase family AAA+ domain containing 2B
PTM: Posttranslational modifications
HBO1: Histone acetyltransferase binding to ORC1
RMSD: Root-mean-square deviation
TSA: Trichostatin A.

## Data availability statement

All the relevant data are contained within this article and in the supporting information.

