## Supporting Information for "Molecular Insights into the Recognition of Acetylated Histone Modifications by the BRPF2 Bromodomain"

**Table of Contents**

1. Supplementary tables S2-S4
2. SDS-PAGE gels and ITC binding plots for the BRPF2 bromodomain S5
3. Histone pull-down and Western blot analysis S6-S8
4. Analysis of mononucleosomes S9
5. Peptide characterization S10-S22

| Peptide | Peptide Sequence | Molecular weight (Da) |
|---|---|---|
| H4unmodified (1-20) | SGRGKGGKGLGKGGAKRHRK | 1992.29 |
| H4K5ac (1-15) | SGRG**Kac**GGKGLGKGGA | 1328.47 |
| H4K8ac (1-15) | SGRGKGG**Kac**GLGKGGA | 1328.47 |
| H4K12ac (1-15) | SGRGKGGKGLG**Kac**GGA | 1328.47 |
| H4K16ac (11-21) | GKGGA**Kac**RHRKVY | 1398.61 |
| H4K5acK8ac (1-15) | SGRG**Kac**GG**Kac**GLGKGGA | 1370.51 |
| H4K5acK12ac (1-15) | SGRG**Kac**GGKGLG**Kac**GGA | 1370.51 |
| H4K8acK12ac (1-15) | SGRGKGG**Kac**GLG**Kac**GGA | 1370.51 |
| H4Kac4 (1-20) | SGRG**Kac**GG**Kac**GLG**Kac**GGA**Kac**RHRK | 2160.44 |
| H3unmodified (1-24) | ARTKQTARKSTGGKAPRKQLATKA | 2554.94 |
| H3K14ac (9-19) | KSTGG**Kac**APRKQ | 1199.36 |
| H2Aunmodified (1-12) | SGRGKQGGKARA | 1172.29 |
| H2AK5ac (1-12) | SGRG**Kac**QGGKARA | 1214.45 |

**Table S1:** List of histone peptides with varying acetylation sites were synthesized and used for this study.

| BRPF2 bromodomain mutants | Primer sequence |
|---|---|
| I586F | GACCCCGCCAGGTTCTTTGCGCAGCCCGTG |
| F587A | GACCCCGCCAGGATAGCCGCGCAGCCCGTG |
| Y599F | GAAGGAGGTACCAGATTTCTTGGATCACATTAAACATCCC |
| N642A | GATAACTGCATGAAGTACGCCGCCAGGGACACCGTGTTC |
| F648A | GCCAGGGACACCGTGGCCTATAGAGCCGCGGTG |

**Table S2:** List of the forward primers designed for site-directed mutagenesis. Reverse primers used are the reverse-complement to the given forward primers.

| Peptide: H4K5ac | Index | Residue | Hydrogen Bonding<br>Distance H-A | Distance D-A | Donor Angle | Donor atom | Acceptor Atom |
|---|---|---|---|---|---|---|---|
| | 1 | K640 | 3.72 | 4.08 | 104.59 | 982 [Ng+] | 646 [O2] |
| | 2 | Y641 | 1.85 | 2.61 | 136.39 | 656 [O3] | 988 [O2] |
| | 3 | Y641 | 3.17 | 3.93 | 134.26 | 970 [Nam] | 656 [O3] |
| | 4 | Y641 | 2.08 | 3.04 | 163.29 | 985 [Nam] | 656 [O3] |
| | 5 | N642 | 1.72 | 2.67 | 160.02 | 664 [Nam] | 997 [O2] |
| | 6 | N642 | 2.25 | 2.87 | 120.23 | 989 [Nam] | 663 [O2] |

| Peptide: H4K5ac | Index | Hydrophobic interactions<br>Residue | Distance | Ligand Atom | Protein Atom |
|---|---|---|---|---|---|
| | 1 | F587 | 3.43 | 998 | 200 |
| | 2 | F648 | 3.45 | 998 | 711 |
| | 3 | F648 | 3.6 | 993 | 712 |
| | 4 | F648 | 3.95 | 991 | 710 |

| Peptide: H4K5ac | Index | Residue | Distance A-W | Water Bridges<br>Distance D-W | Donor Angle | Water Angle | Donor Atom | Acceptor Atom | Water Atom |
|---|---|---|---|---|---|---|---|---|---|
| | 1 | K594 | 2.87 | 3.48 | 107.79 | 96.85 | 964 [O3] | 255 [O2] | 33316 |
| | 2 | E595 | 3.2 | 4.01 | 149.13 | 74.2 | 1001 [Nam] | 264 [O2] | 27954 |
| | 3 | Y599 | 3.8 | 2.6 | 174.3 | 75.07 | 296 [O3] | 995 [Nam] | 28764 |
| | 4 | H602 | 3.61 | 3.67 | 126.41 | 101.66 | 981 [Ng+] | 324 [O2] | 34147 |
| | 5 | H602 | 2.74 | 3.81 | 161.2 | 82.79 | 981 [Ng+] | 324 [O2] | 29765 |
| | 6 | H602 | 3.78 | 3.23 | 118.33 | 115.21 | 319 [Nar] | 981 [Ng+] | 34038 |
| | 7 | Y641 | 4.06 | 3.33 | 160.24 | 126.99 | 974 [Nam] | 656 [O3] | 33942 |
| | 8 | A643 | 4.01 | 2.87 | 177.58 | 110.64 | 667 [Nam] | 984 [O2] | 32986 |

**Table S3:** Summary of hydrogen bonds, hydrophobic interactions and water bridges between the H4K5ac peptide and the BRPF2 bromodomain.

| Peptide: H4K5acK12ac | Index | Residue | Hydrogen Bonding Distance H-A | Distance D-A | Donor Angle | Donor atom | Acceptor Atom |
|---|---|---|---|---|---|---|---|
| | 1 | K594 | 2.65 | 3.26 | 118.13 | 1015 [N3] | 255 [O2] |
| | 2 | E595 | 2.54 | 3.22 | 126.69 | 1001 [Nam] | 262 [O3] |
| | 3 | E595 | 3.55 | 3.85 | 101.84 | 262 [O3] | 1004 [O2] |
| | 4 | Y641 | 1.86 | 2.71 | 150.09 | 656 [O3] | 988 [O2] |
| | 5 | Y641 | 2.21 | 3.14 | 156.23 | 985 [Nam] | 656 [O3] |
| | 6 | Y641 | 2.83 | 3.52 | 128.25 | 970 [Nam] | 656 [O3] |
| | 7 | Y641 | 2.5 | 2.92 | 105.8 | 974 [Nam] | 656 [O3] |
| | 8 | N642 | 3.1 | 4.04 | 159.5 | 664 [Nam] | 995 [Nam] |
| | 9 | A643 | 1.96 | 2.93 | 168.92 | 667 [Nam] | 984 [O2] |

| Peptide: H4K5acK12ac | Index | Residue | Hydrophobic Interactions Distance | Ligand Atom | Protein Atom |
|---|---|---|---|---|---|
| | 1 | A643 | 3.62 | 977 | 669 |
| | 2 | F648 | 3.55 | 993 | 713 |

| Peptide: H4K5acK12ac | Index | Residue | Water Bridges Distance A-W | Distance D-W | Donor Angle | Water Angle | Donor Atom | AcceptorAtom | Water Atom |
|---|---|---|---|---|---|---|---|---|---|
| | 1 | I586 | 4.04 | 3.39 | 130.35 | 95.33 | 995 [Nam] | 193 [O2] | 27780 |
| | 2 | E595 | 3.44 | 3.78 | 117.85 | 77.36 | 967 [N3] | 264 [O2] | 33437 |
| | 3 | Y599 | 2.74 | 2.71 | 169.71 | 75.58 | 296 [O3] | 997 [O2] | 28766 |
| | 4 | Y641 | 3.69 | 3.47 | 165.53 | 71.61 | 967 [N3] | 656 [O3] | 36233 |
| | 5 | Y641 | 3.96 | 3.99 | 145.72 | 79.81 | 982 [Ng+] | 658 [O2] | 34810 |

**Table S4:** Summary of hydrogen bonds, hydrophobic interactions and water bridges between the H4K5acK12ac peptide and the BRPF2 bromodomain.

| BRPF2 bromodomain | Molar ellipticity at 222 nm | % α-helix | % β-sheet |
|---|---|---|---|
| WT | -1114570 | 91.55 | 0.49 |
| I586F | -949206 | 89.55 | 0.5 |
| F587A | -974544 | 90.54 | 0.5 |
| Y599F | -1026746 | 91.54 | 0.49 |
| N642A | -1070848 | 91.54 | 0.49 |
| F648A | -949206 | 89.55 | 0.5 |

**Table S5:** The molar ellipticity at 222 nm and the relative percentage of α-helix and β-sheet composition for the wild-type BRPF2 bromodomain and its mutants, calculated from circular dichroism experiments.

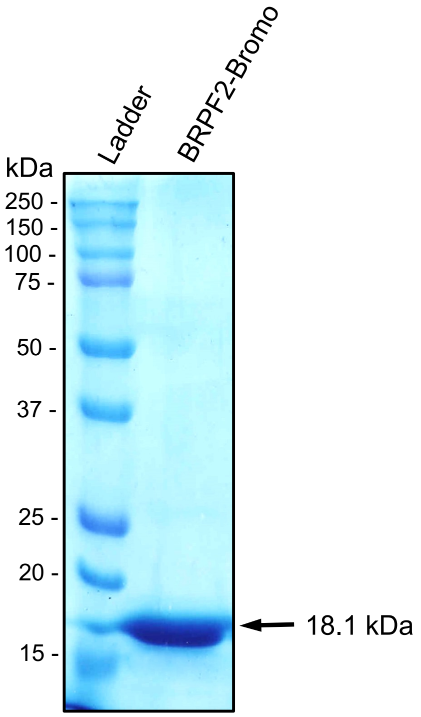


**Figure S1:** Coomassie blue staining of gel showing the purity of the overexpressed wild-type BRPF2 bromodomain.

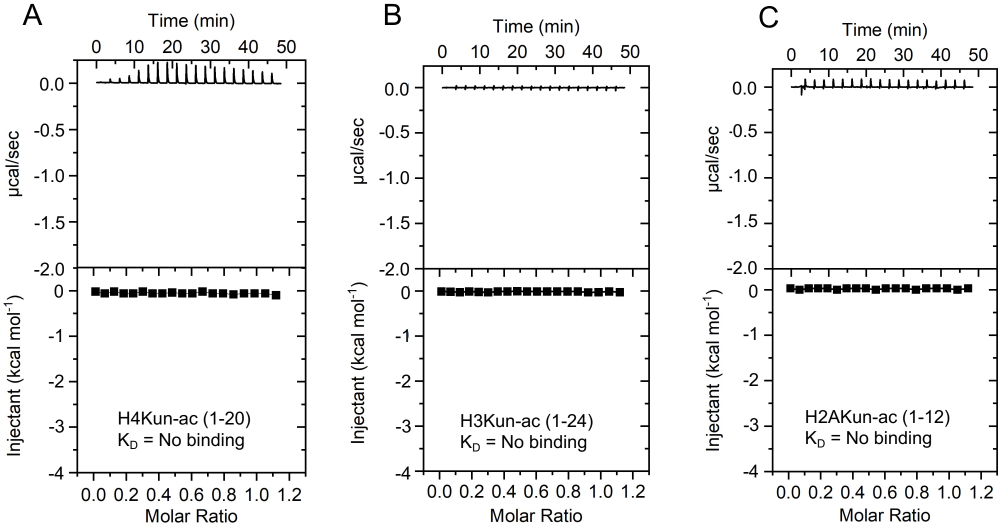


**Figure S2:** (A-C) ITC binding plots for the interaction of BRPF2 bromodomain with unacetylated H4, H3, and H2A peptides.

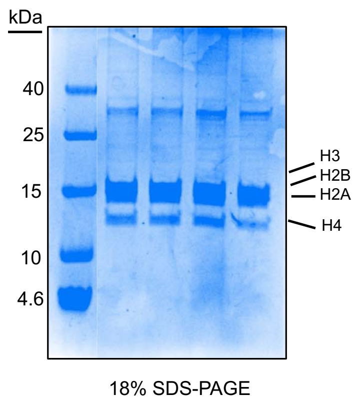


**Figure S3:** Acid extracted endogenous histones from HeLa cells were run on a 18% SDS-PAGE gel and stained with Coomassie blue dye.

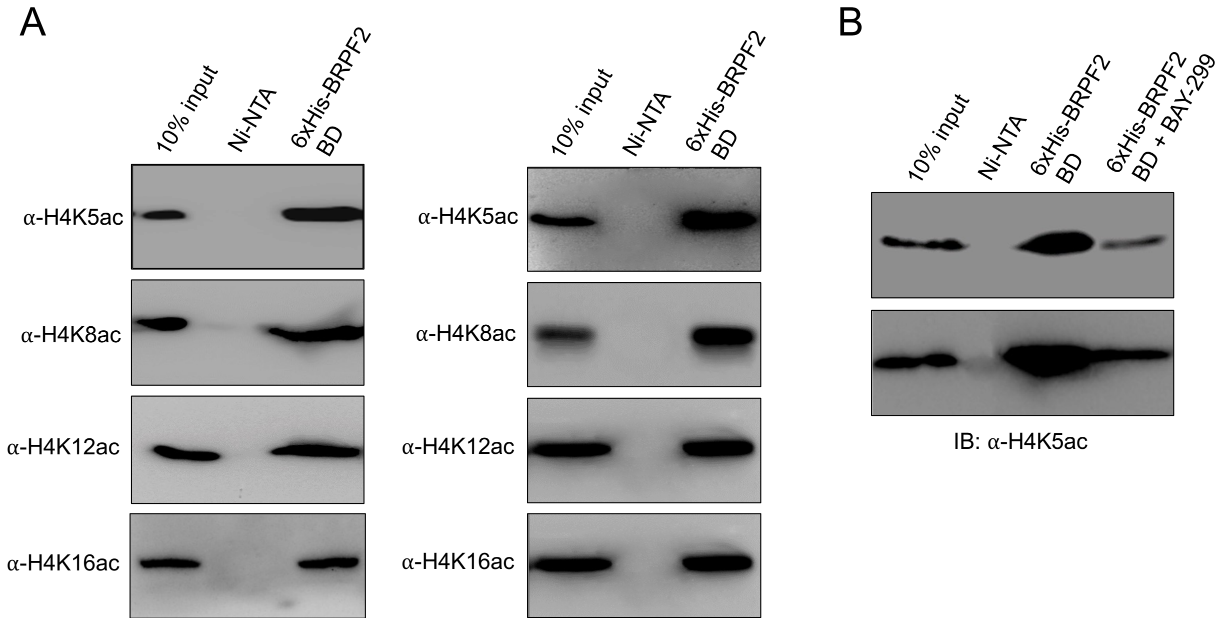


**Figure S4:** Interaction of BRPF2 bromodomain with endogenous histone H4. (A) The BRPF2 bromodomain was incubated with hyperacetylated H4 and bound acetylated H4 was enriched by Ni-NTA bead, followed by western blot against anti-acetylated H4 antibodies (biological replicates). (B) The BRPF2 bromodomain was incubated with hyperacetylated H4 in the presence or absence of BAY-299 inhibitor, and bound acetylated H4 was enriched by Ni-NTA bead, followed by western blot against anti-H4K5ac antibody (biological replicates).

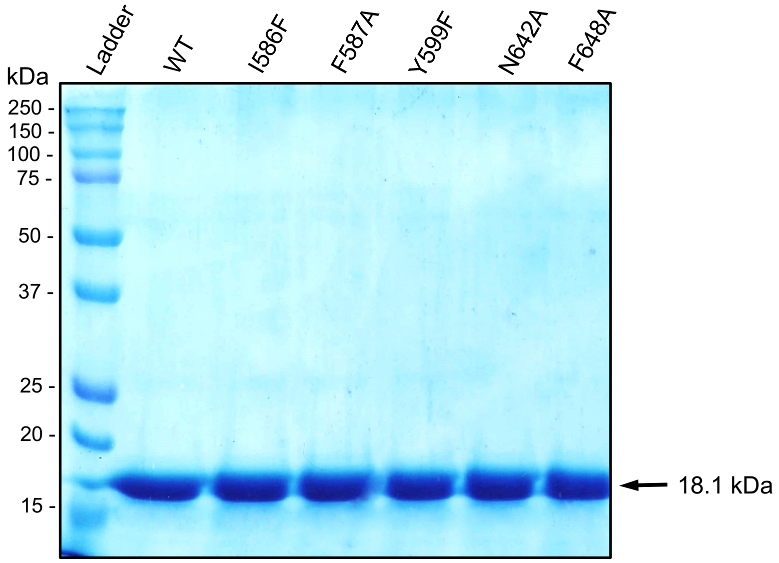


**Figure S5:** Coomassie blue staining of gel showing the protein expression and purity of wild-type BRPF2 bromodomain and its mutant proteins.

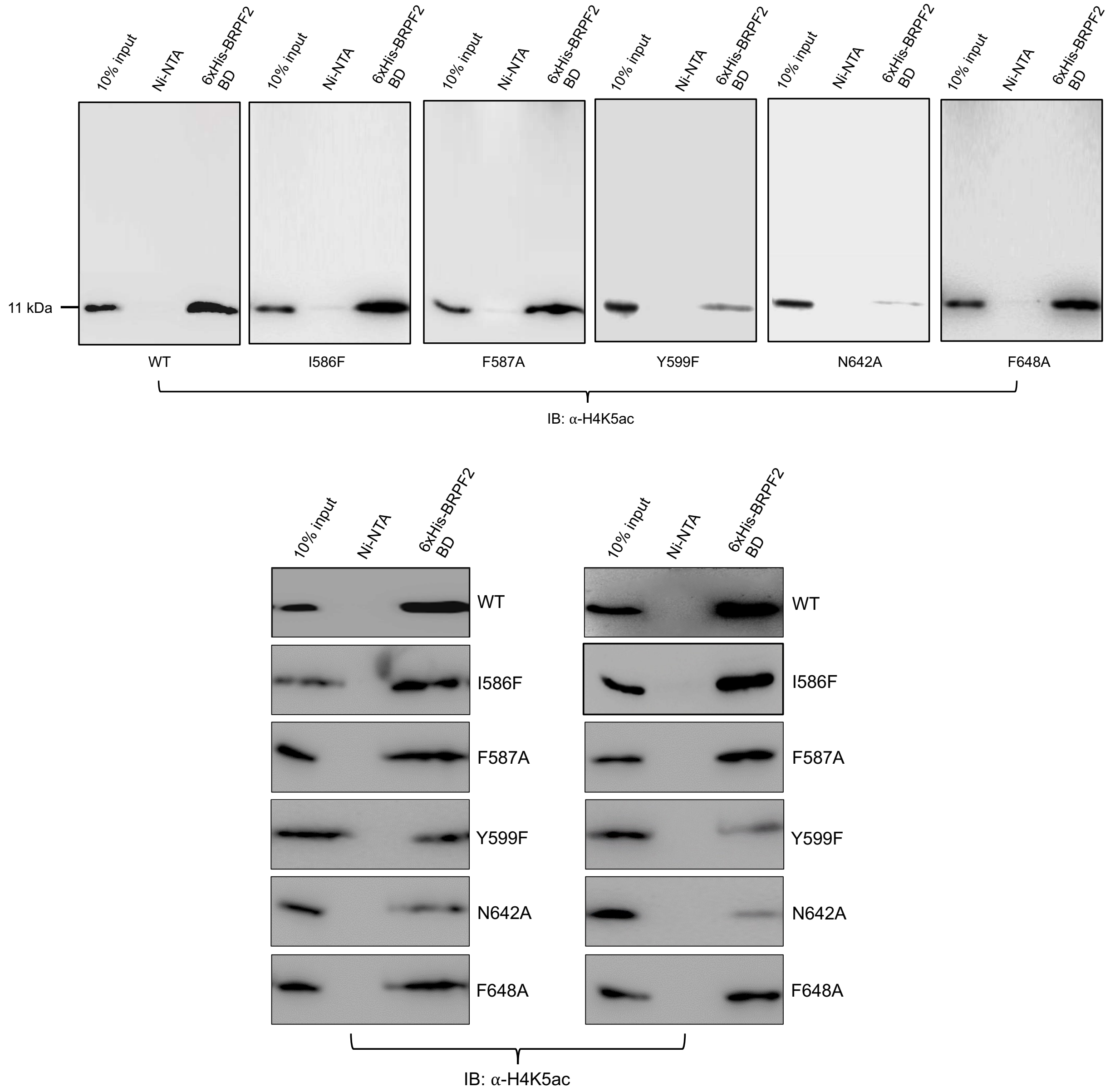


**Figure S6:** Interaction of BRPF2 bromodomain wild-type and its mutants with endogenous histone H4. The mutational analysis of acetyllysine binding pocket residues demonstrates that the Y599F and N642A mutants significantly abolished the interaction of BRPF2 bromodomain with H4K5ac modification as compared to the other mutants (biological replicates).

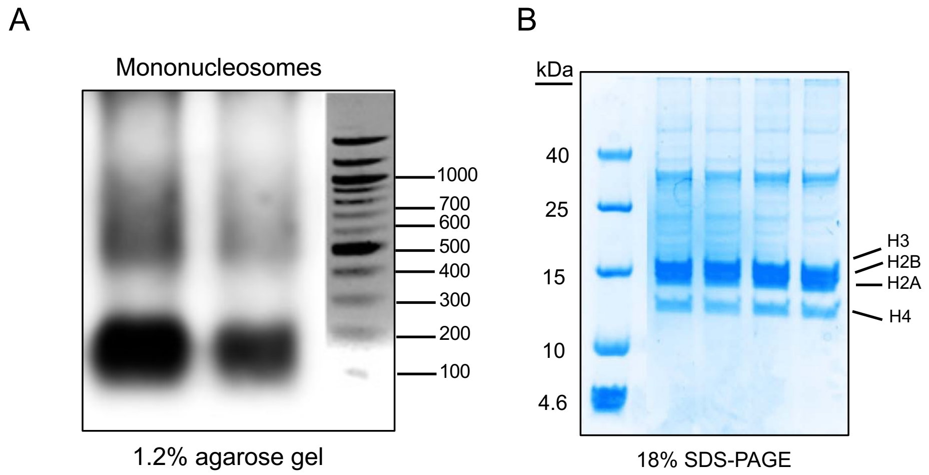


**Figure S7:** Mononucleosomes were isolated from HeLa cells. (A) The mononucleosomal pool analyzed on 1.2% agarose gel. (B) Presence of histones in the isolated mononucleosomal pool.

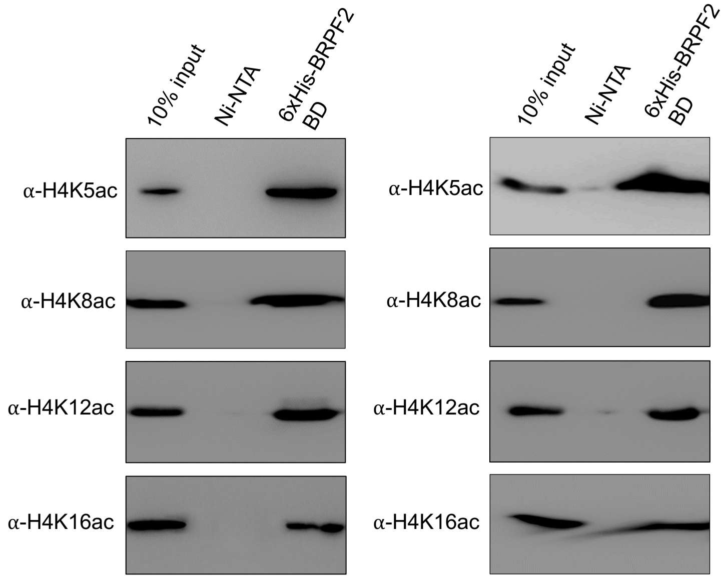


**Figure S8:** The BRPF2 bromodomain strongly recognizes the mono-acetylated H4K5ac mark at the mononucleosome level. Recombinantly produced 6xHis-tagged BRPF2 bromodomain was used to pull-down native mononucleosomes, and the bound material was probed by western blot against the anti-acetylated H4 antibodies (biological replicates).

**H4 unmodified (1-20)**

**HPLC REPORT**

Sample: Pep-175 SGRGKGGKGLGKGGAKRHRK — Analyzed date: 27-09-2020
Analyst: Dr.RS-SBio
Column: Symmetrix ODS-R, 4.6*250mm, 5μm
Solvent A: A: 0.1% Trifluoroacetic Acid in 100% Acetonitrile
Solvent B: B: 0.1% Trifluoroacetic Acid in 100% Water

Gradient:

| | A | B |
|---|---|---|
| 0.0min | 9% | 91% |
| 25.0min | 34% | 56% |
| 25.1min | 100% | 0% |
| 30.0min | Stop | |

Volume: 20μl
Wavelength: 220nm
Flow rate: 1.0ml/min

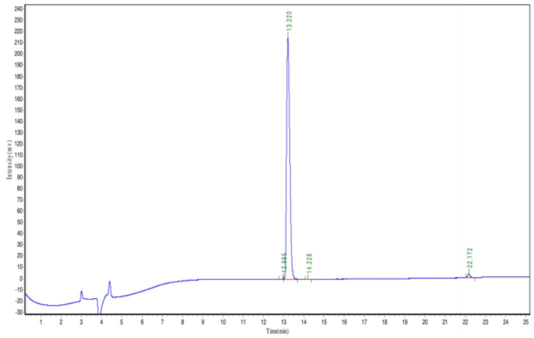


| Peak | Time | Height | Area | Conc. |
|---|---|---|---|---|
| 1 | 12.985 | 285.682 | 1626.532 | 0.0688 |
| 2 | 13.220 | 215636.578 | 2333620.000 | 98.7647 |
| 3 | 14.228 | 193.103 | 1689.600 | 0.0715 |
| 4 | 22.172 | 3093.963 | 25871.998 | 1.0950 |
| Total | | | | 100.000 |

**Figure S9:** HPLC purity trace for the H4 unmodified (1-20) peptide.

**H4 unmodified (1-20)**

**MASS SPECTROMETRY REPORT**

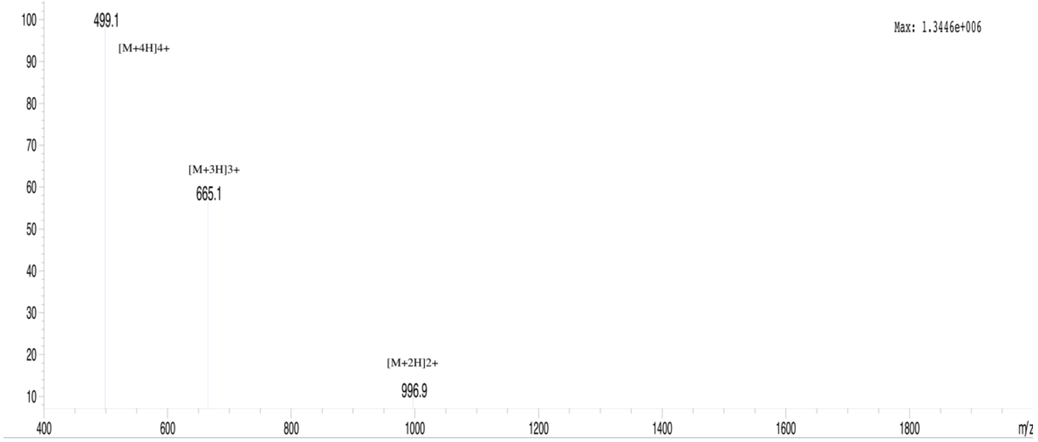


| Sample Description | | Instrument | Agilent-6125B | | |
|---|---|---|---|---|---|
| Analyzed date: | 27-09-2020 | Probe: | ESI | Probe Bias: | +4.5kv |
| Analyst: | Dr.AR-SBio | Nebulizer Gas Flow: | 1.5L/min | Detector: | 1.5kv |
| Sample: | Pep-175 SGRGKGGKGLGKGGAKRHRK | CDL: | -20.0v | T. Flow: | 0.2ml/min |
| M.W.: | 1992.29 | CDL Temp.: | 250 °C | B. Conc.: | 50%H2O/50%ACN |

**Figure S10:** MS spectra for the H4 unmodified (1-20) peptide.

**HPLC REPORT**

**H4K5ac (1-15)**

Sample: Pep-458 SGRGK(ac)GGKGLGKGGA Analyzed date: 19-03-2022
Analyst: Dr.RS-SBio
Column: 4.6×250mm,ChromCore 120 C18 5um
Solvent A A: 0.1% Trifluoroacetic Acid in 100% Acetonitrile
Solvent B B: 0.1% Trifluoroacetic Acid in 100% Water
Gradient:

| | A | B |
|---|---|---|
| 0.0min | 10% | 90% |
| 25.0min | 35% | 65% |
| 25.1min | 100% | 0% |
| 30.0min | Stop | |

Volume: 10μl
Wavelength: 220nm
Flow rate: 1.0ml/min

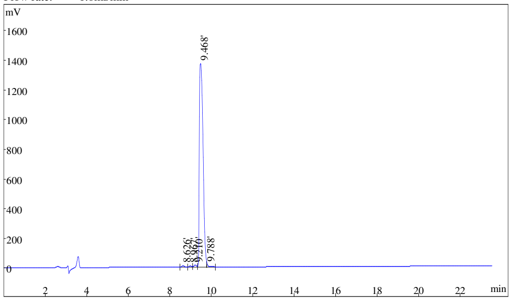


| Rank | Time | Quantity | Area | Height |
|---|---|---|---|---|
| 1 | 8.626 | 0.3913 | 62159 | 7970 |
| 2 | 8.967 | 0.3271 | 51969 | 6061 |
| 3 | 9.210 | 0.6437 | 102265 | 13485 |
| 4 | 9.468 | 98.14 | 15590126 | 1372133 |
| 5 | 9.788 | 0.502 | 79753 | 16541 |
| Total | | 100 | 15886272 | 1416190 |

**Figure S11:** HPLC purity trace for the H4K5ac (1-15) peptide.

**H4K5ac (1-15)**

**MASS SPECTROMETRY REPORT**

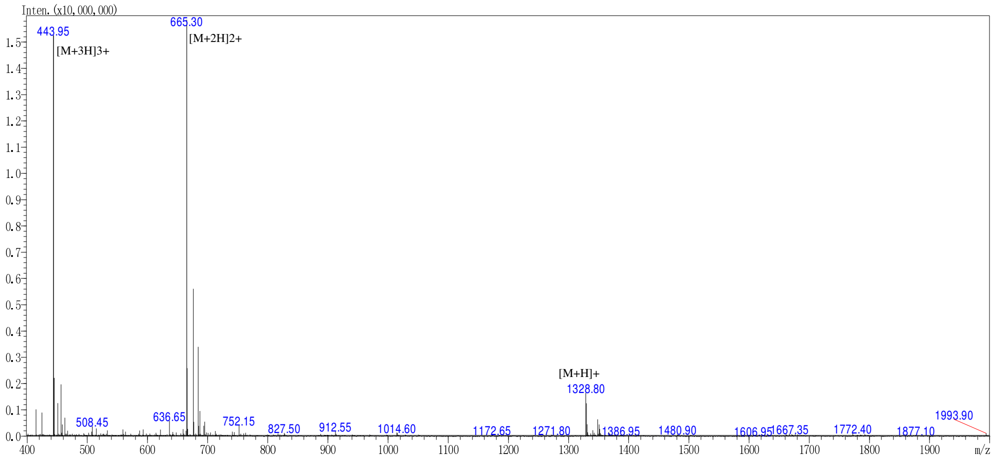


| Sample Description | | Instrument | SHIMADZU LCMS-2020 | | |
|---|---|---|---|---|---|
| Analyzed date: | 20-03-2022 | Probe: | ESI | Probe Bias: | +4.5kv |
| Analyst: | Dr.AR-SBio | Nebulizer Gas Flow: | 1.5L/min | Detector: | 1.2kv |
| Sample: | Pep-458 SGRGK(ac)GGKGLGKGGA | CDL: | -20.0v | T. Flow: | 0.2ml/min |
| M.W.: | 1328.47 | CDL Temp.: | 250 °C | B. Conc.: | 50%H2O/50%ACN |
| Lot. No.: | P220222-MX974607 | Block Temp.: | 400 °C | | |

**Figure S12:** MS spectra for the H4K5ac (1-15) peptide.

### HPLC REPORT
## H4K8ac (1-15)

Sample: Pep-173 SGRGKGGK(ac)GLGKGGA    Analyzed date: 26-09-2020
Analyst: Dr.RS-SBio
Column: Symmetrix ODS-R, 4.6*250mm, 5μm
Solvent A: A: 0.1% Trifluoroacetic Acid in 100% Acetonitrile
Solvent B: B: 0.1% Trifluoroacetic Acid in 100% Water
Gradient:

| | A | B |
|---|---|---|
| 0.0min | 2% | 98% |
| 25.0min | 27% | 73% |
| 25.1min | 100% | 0% |
| 30.0min | Stop | |

Volume: 20μl
Wavelength: 220nm
Flow rate: 1.0ml/min

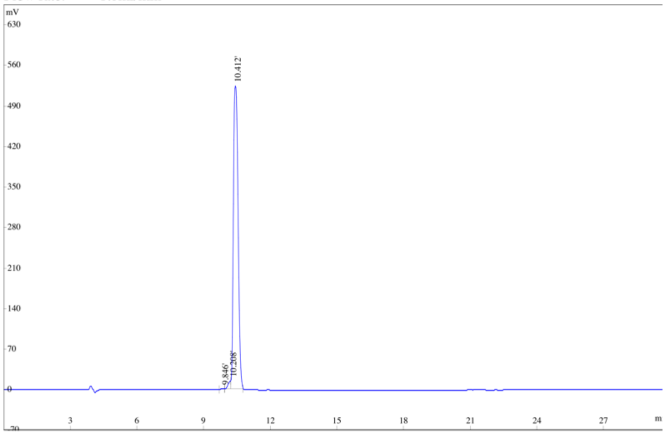


| Rank | Time | Conc. | Area | Height |
|---|---|---|---|---|
| 1 | 9.846 | 0.0795 | 5629 | 838 |
| 2 | 10.208 | 1.5212 | 107725 | 13486 |
| 3 | 10.412 | 98.3993 | 6968244 | 521870 |
| Total | | 100 | 7081598 | 536194 |

**Figure S13:** HPLC purity trace for the H4K8ac (1-15) peptide.

### H4K8ac (1-15)
#### MASS SPECTROMETRY REPORT

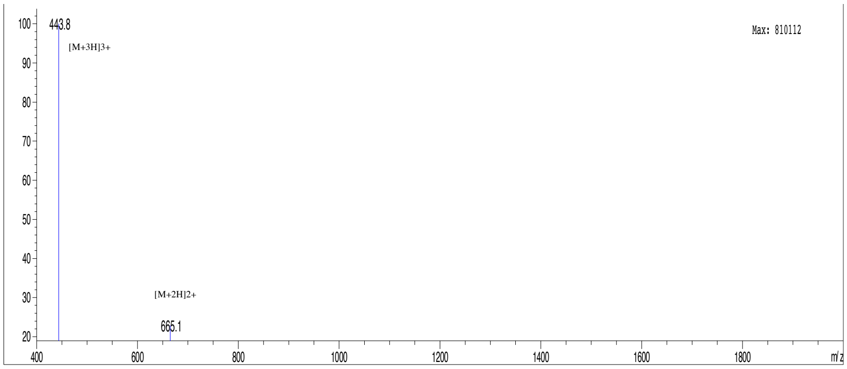


Sample Description

| | | | | | |
|---|---|---|---|---|---|
| Analyzed date: | 28-09-2020 | Instrument | Agilent-6125B | | |
| Analyst: | Dr.AR-Sbio | Probe: | ESI | Probe Bias: | +4.5kv |
| Sample: | Pep-173 SGRGKGGK(ac)GLGKGGA | Nebulizer Gas Flow: | 1.5L/min | Detector: | 1.5kv |
| M.W.: | 1328.47 | CDL: | -20.0v | T. Flow: | 0.2ml/min |
| | | CDL Temp.: | 250 °C | B. Conc.: | 50%H2O/50%ACN |

**Figure S14:** MS spectra for the H4K8ac (1-15) peptide.

**HPLC REPORT**

**H4K12ac (1-15)**

| | | | |
|---|---|---|---|
| Sample: | Pep-459 SGRGKGGKGLGK(ac)GGA | | Analyzed date: 18-03-2022 |
| Analyst: | Dr.RS-SBio | | |
| Column: | 4.6×250mm,ChromCore 120 C18 5um | | |
| Solvent A | A: 0.1% Trifluoroacetic Acid in 100% Acetonitrile | | |
| Solvent B | B: 0.1% Trifluoroacetic Acid in 100% Water | | |
| Gradient: | | A | B |
| | 0.0min | 10% | 90% |
| | 25.0min | 35% | 65% |
| | 25.1min | 100% | 0% |
| | 30.0min | Stop | |
| Volume: | 10µl | | |
| Wavelength: | 220nm | | |
| Flow rate: | 1.0ml/min | | |

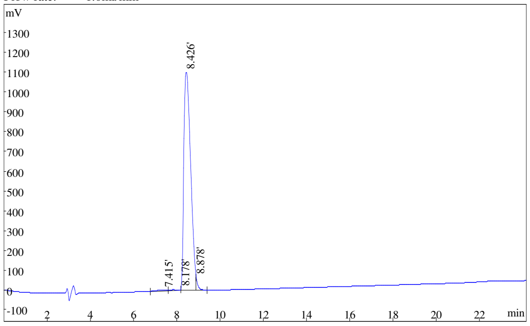


| Rank | Time | Conc. | Area | Height |
|---|---|---|---|---|
| 1 | 7.415 | 0.7537 | 180164 | 7551 |
| 2 | 8.178 | 0.6391 | 152758 | 7811 |
| 3 | 8.426 | 97.23 | 23239382 | 1100502 |
| 4 | 8.878 | 1.381 | 330166 | 64707 |
| Total | | 100 | 23902470 | 1180571 |

**Figure S15:** HPLC purity trace for the H4K12ac (1-15) peptide.

**H4K12ac (1-15)**

**MASS SPECTROMETRY REPORT**

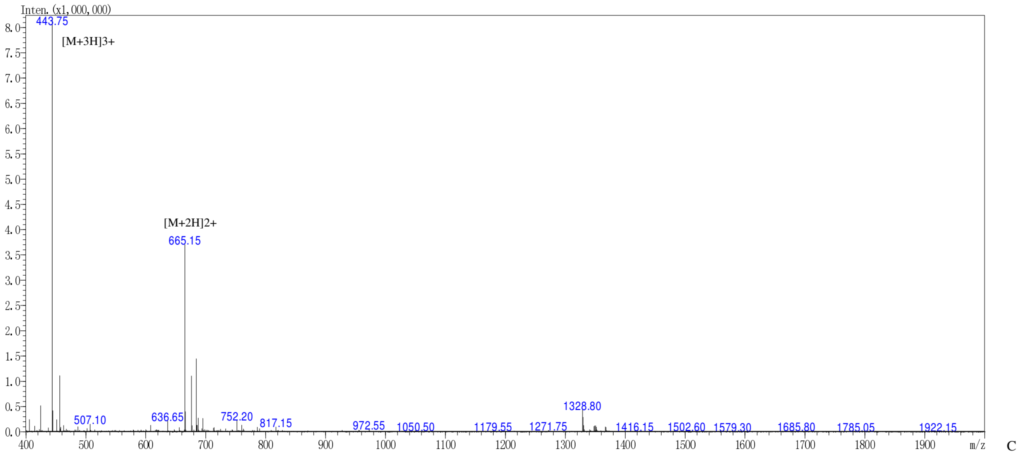


| | | | | | |
|---|---|---|---|---|---|
| Sample Description | | Instrument | SHIMADZU LCMS-2020 | | |
| Analyzed date: | 19-03-2022 | Probe: | ESI | Probe Bias: | +4.5kv |
| Analyst: | Dr.AR-SBio | Nebulizer Gas Flow: | 1.5L/min | Detector: | 1.2kv |
| Sample: | Pep-459 SGRGKGGKGLGK(ac)GGA | CDL: | -20.0v | T. Flow: | 0.2ml/min |
| M.W.: | 1328.47 | CDL Temp.: | 250 °C | B. Conc.: | 50%H2O/50%ACN |

**Figure S16:** MS spectra for the H4K12ac (1-15) peptide.

**H4K5acK8ac (1-15)**

### HPLC REPORT

Sample: Pep-460 SGRGK(ac)GGK(ac)GLGKGGA Analyzed date: 21-03-2022
Analyst: Dr.RS-SBio
Column: 4.6×250mm,ChromCore 120 C18 5um
Solvent A A: 0.1% Trifluoroacetic Acid in 100% Acetonitrile
Solvent B B: 0.1% Trifluoroacetic Acid in 100% Water
Gradient:

| | A | B |
|---|---|---|
| 0.0min | 10% | 90% |
| 25.0min | 35% | 65% |
| 25.1min | 100% | 0% |
| 30.0min | Stop | |

Volume: 10µl
Wavelength: 220nm
Flow rate: 1.0ml/min

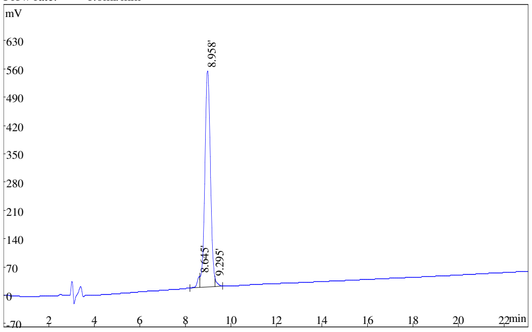


| Rank | Time | Conc. | Area | Height |
|---|---|---|---|---|
| 1 | 8.645 | 2.669 | 238653 | 27779 |
| 2 | 8.958 | 96.19 | 8601225 | 532768 |
| 3 | 9.295 | 1.145 | 102427 | 17006 |
| Total | | 100 | 8942305 | 577553 |

**Figure S17:** HPLC purity trace for the H4K5acK8ac (1-15) peptide.

**H4K5acK8ac (1-15)**
**MASS SPECTROMETRY REPORT**

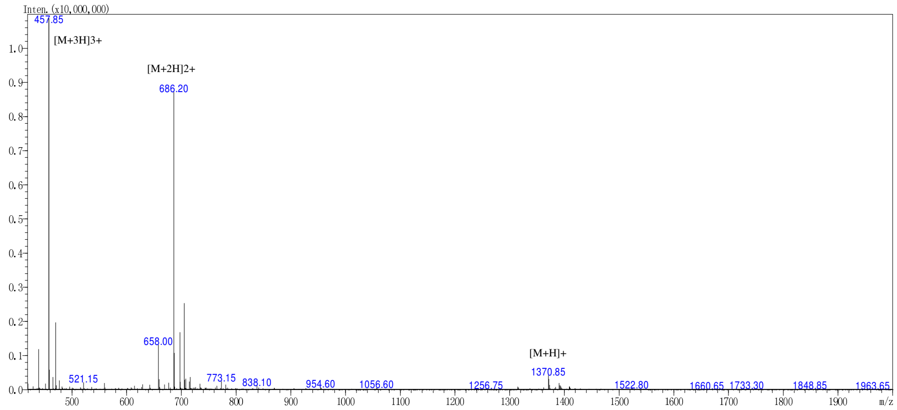


| | | | | | |
|---|---|---|---|---|---|
| Sample Description | | Instrument | SHIMADZU LCMS-2020 | | |
| Analyzed date: | 22-03-2022 | Probe: | ESI | Probe Bias: | +4.5kv |
| Analyst: | Dr.RS-SBio | Nebulizer Gas Flow: | 1.5L/min | Detector: | 1.2kv |
| Sample: | Pep-460 SGRGK(ac)GGK(ac)GLGKGGA | CDL: | -20.0v | T. Flow: | 0.2ml/min |
| M.W.: | 1370.51 | CDL Temp.: | 250 °C | B. Conc.: | 50%H2O/50%ACN |

**Figure S18:** MS spectra for the H4K5acK8ac (1-15) peptide.

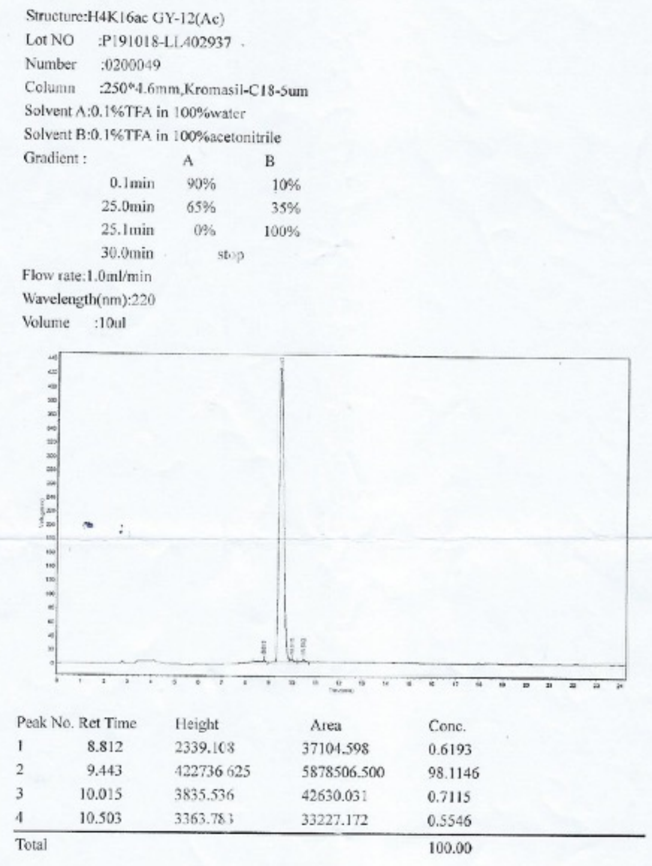

Structure:H4K16ac GY-12(Ac)
Lot NO :P191018-LL402937
Number :0200049
Column :250*4.6mm,Kromasil-C18-5um
Solvent A:0.1%TFA in 100%water
Solvent B:0.1%TFA in 100%acetonitrile
Gradient :

| | A | B |
|---|---|---|
| 0.1min | 90% | 10% |
| 25.0min | 65% | 35% |
| 25.1min | 0% | 100% |
| 30.0min | stop | |

Flow rate:1.0ml/min
Wavelength(nm):220
Volume :10ul

| Peak No. | Ret Time | Height | Area | Conc. |
|---|---|---|---|---|
| 1 | 8.812 | 2339.108 | 37104.598 | 0.6193 |
| 2 | 9.443 | 422736.625 | 5878506.500 | 98.1146 |
| 3 | 10.015 | 3835.536 | 42630.031 | 0.7115 |
| 4 | 10.503 | 3363.783 | 33227.172 | 0.5546 |
| Total | | | | 100.00 |

**Figure S19:** HPLC purity trace for the H4K16ac (11-21) peptide.

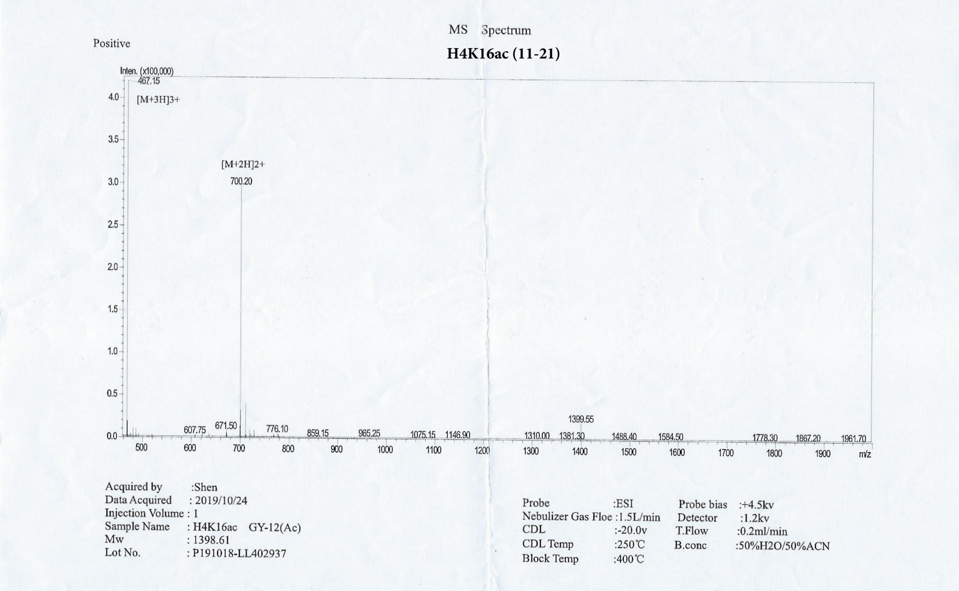


**Figure S20:** MS spectra for the H4K16ac (11-21) peptide.

**H4K5acK12ac (1-15)**

**HPLC REPORT**

Sample: Pep-460 SGRGK(ac)GGK(ac)GLGKGGA Analyzed date: 21-03-2022
Analyst: Dr.RS-SBio
Column: 4.6×250mm,ChromCore 120 C18 5um
Solvent A A: 0.1% Trifluoroacetic Acid in 100% Acetonitrile
Solvent B B: 0.1% Trifluoroacetic Acid in 100% Water
Gradient:

| | A | B |
|---|---|---|
| 0.0min | 10% | 90% |
| 25.0min | 35% | 65% |
| 25.1min | 100% | 0% |
| 30.0min | Stop | |

Volume: 10μl
Wavelength: 220nm
Flow rate: 1.0ml/min

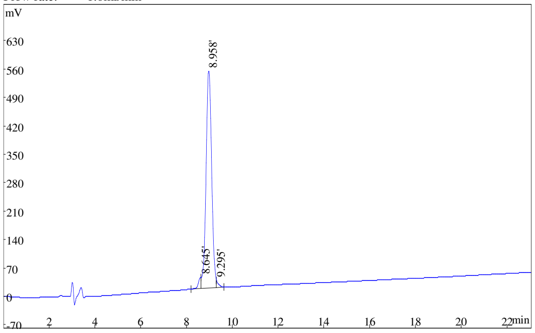


| Rank | Time | Conc. | Area | Height |
|---|---|---|---|---|
| 1 | 8.645 | 2.669 | 238653 | 27779 |
| 2 | 8.958 | 96.19 | 8601225 | 532768 |
| 3 | 9.295 | 1.145 | 102427 | 17006 |
| Total | | 100 | 8942305 | 577553 |

**Figure S21:** HPLC purity trace for the H4K5acK12ac (1-15) peptide.

**H4K5acK12ac (1-15)**

**MASS SPECTROMETRY REPORT**

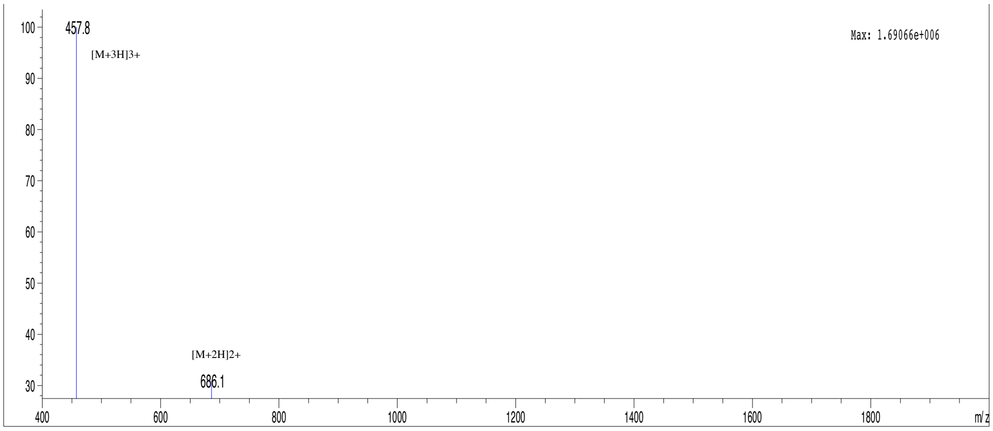


| Sample Description | | Instrument | Agilent-6125B | | |
|---|---|---|---|---|---|
| Analyzed date: | 03-10-2020 | Probe: | ESI | Probe Bias: | +4.5kv |
| Analyst: | Dr.AR-SBio | Nebulizer Gas Flow: | 1.5L/min | Detector: | 1.5kv |
| Sample: | Pep-171 SGRGK(ac)GGKGLGK(ac)GGA | CDL: | -20.0v | T. Flow: | 0.2ml/min |
| M.W.: | 1370.51 | CDL Temp.: | 250 °C | B. Conc.: | 50%H2O/50%ACN |

**Figure S22:** MS spectra for the H4K5acK12ac (1-15) peptide.

**H4K8acK12ac**

### HPLC REPORT

| | | | |
|---|---|---|---|
| Sample: | Pep-461 SGRGKGGK(ac)GLGK(ac)GGA | | Analyzed date: 20-03-2022 |
| Analyst: | Dr.RS-SBio | | |
| Column: | Kromasil-C18, 4.6*250mm, 5µm | | |
| Solvent A | A: 0.1% Trifluoroacetic Acid in 100% Acetonitrile | | |
| Solvent B | B: 0.1% Trifluoroacetic Acid in 100% Water | | |
| Gradient: | | A | B |
| | 0.0min | 9% | 91% |
| | 25.0min | 34% | 66% |
| | 25.1min | 100% | 0% |
| | 30.0min | Stop | |
| Volume: | 10µl | | |
| Wavelength: | 220nm | | |
| Flow rate: | 1.0ml/min | | |

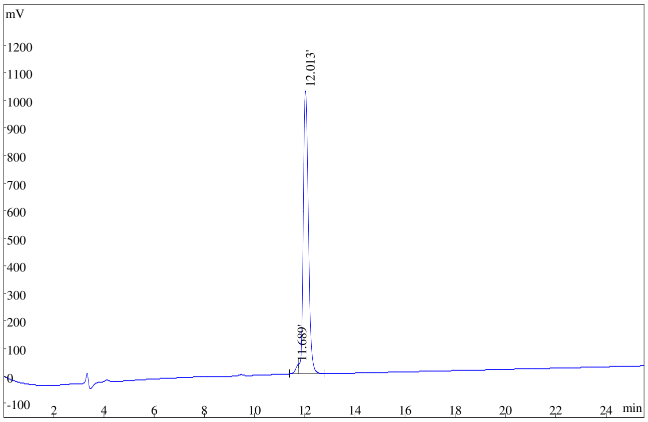


| Rank | Time | Area | Conc. |
|---|---|---|---|
| 1 | 11.689 | 271397 | 1.976 |
| 2 | 12.013 | 13463240 | 98.02 |
| Total | | 13734637 | 100 |

**Figure S23:** HPLC purity trace for the H4K8acK12ac (1-15) peptide.

**H4K8acK12ac (1-15)**

**MASS SPECTROMETRY REPORT**

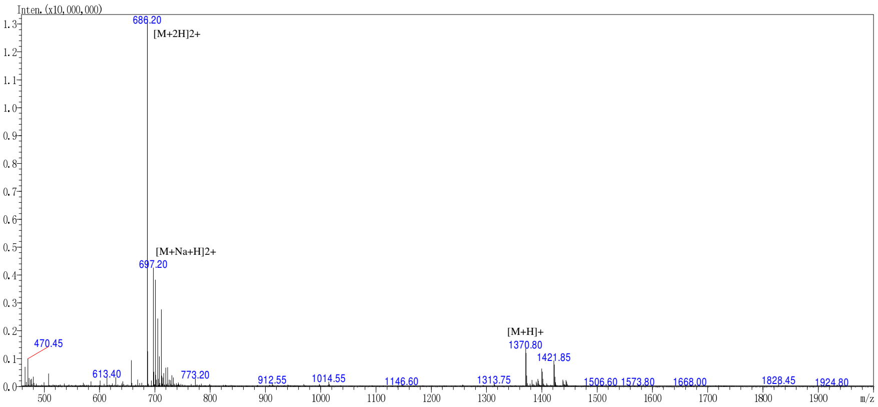


| | | | | | |
|---|---|---|---|---|---|
| Sample Description | | Instrument | SHIMADZU LCMS-2020 | | |
| Analyzed date: | 21-03-2022 | Probe: | ESI | Probe Bias: | +4.5kv |
| Analyst: | Dr.AR-SBio | Nebulizer Gas Flow: | 1.5L/min | Detector: | 1.2kv |
| Sample: | Pep-461 SGRGKGGK(ac)GLGK(ac)GGA | CDL: | -20.0v | T. Flow: | 0.2ml/min |
| M.W.: | 1370.51 | CDL Temp.: | 250 °C | B. Conc.: | 50%H2O/50%ACN |

**Figure S24:** MS spectra for the H4K8acK12ac (1-15) peptide.

**HPLC REPORT**
**H4Kac4 (1-20)**

| | | |
|---|---|---|
| Sample: | Pep-197 SGRGK(ac)GGK(ac)GLGK(ac)GGAK(ac)RHRK | Analyzed date: 11-12-2020 |
| Analyst: | Dr.RS-SBio | |
| Column: | Gemini-NX 5μ C18 110A, 4.6*250mm | |
| Solvent A | A: 0.1% Trifluoroacetic Acid in 100% Acetonitrile | |
| Solvent B | B: 0.1% Trifluoroacetic Acid in 100% Water | |

Gradient:

| | A | B |
|---|---|---|
| 0.0min | 10% | 90% |
| 25.0min | 35% | 65% |
| 25.1min | 100% | 0% |
| 30.0min | Stop | |

Volume: 20μl
Wavelength: 220nm
Flow rate: 1.0ml/min

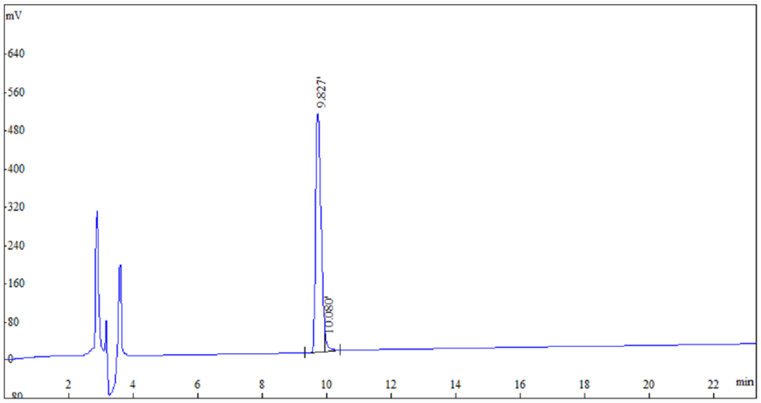


| Rank | Time | Conc. | Area | Height |
|---|---|---|---|---|
| 1 | 9.827 | 97.5303 | 5575313 | 499399 |
| 2 | 10.080 | 2.4697 | 141182 | 25521 |
| Total | | 100 | 5716495 | 524920 |

**Figure S25:** HPLC purity trace for the H4Kac4 (1-20) peptide.

**MASS SPECTROMETRY REPORT**

H4Kac4 (1-20)

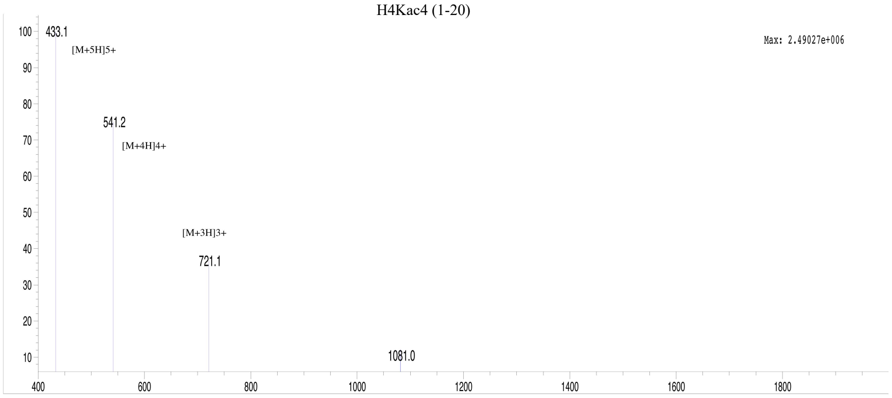


| | | | | | |
|---|---|---|---|---|---|
| Sample Description | | Instrument | Agilent-6125B | | |
| Analyzed date: | 12-12-2020 | Probe: | ESI | Probe Bias: | 4.5kv |
| Analyst: | Dr.AR-SBio | Nebulizer Gas Flow: | 1.5L/min | Detector: | 1.5kv |
| Sample: | Pep-197 SGRGK(ac)GGK(ac)GLGK(ac)GGAK(ac)RHRK | CDL: | -20.0v | T. Flow: | 0.2ml/min |
| M.W.: | 2160.44 | CDL Temp.: | 250 °C | B. Conc.: | 50%H2O/50%ACN |

**Figure S26:** MS spectra for the H4Kac4 (1-20) peptide.

**HPLC REPORT**

**H2A unmodified (1-12)**

Sample: Pep-177 SGRGKQGGKARA — Analyzed date: 24-09-2020

Analyst: Dr.RS-SBio

Column: 4.6×250mm,Sinochrom ODS-BP 5μm

Solvent A: A: 0.1% Trifluoroacetic Acid in 100% Acetonitrile

Solvent B: B: 0.1% Trifluoroacetic Acid in 100% Water

Gradient:

| | A | B |
|---|---|---|
| 0.0min | 2% | 98% |
| 25.0min | 27% | 73% |
| 25.1min | 100% | 0% |
| 30.0min | | Stop |

Volume: 5μl

Wavelength: 220nm

Flow rate: 1.0ml/min

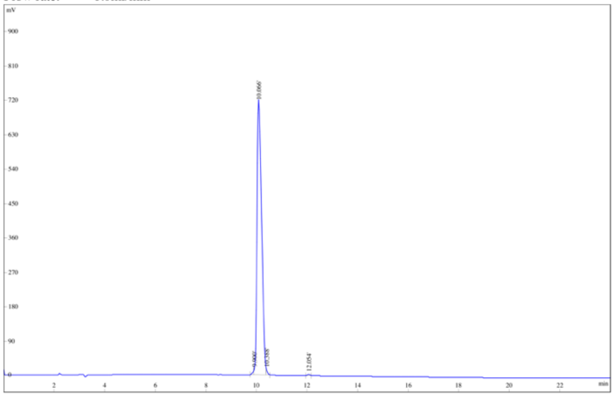


| Rank | Time | Conc. | Area | Height |
|---|---|---|---|---|
| 1 | 9.900 | 0.5689 | 53051 | 17056 |
| 2 | 10.066 | 98.89 | 9222006 | 723039 |
| 3 | 10.388 | 0.327 | 30492 | 11547 |
| 4 | 12.054 | 0.2133 | 19896 | 2864 |
| Total | | 100 | 9325445 | 754506 |

**Figure S27:** HPLC purity trace for the H2A unmodified (1-12) peptide.

**H2A unmodified (1-12)**

**MASS SPECTROMETRY REPORT**

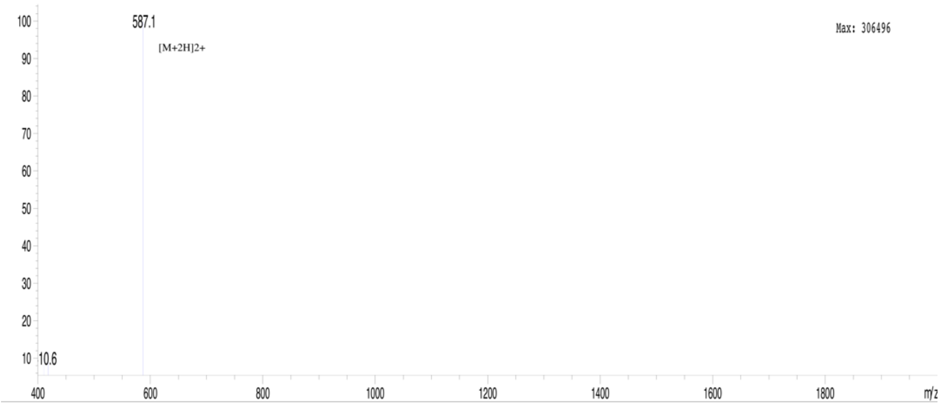


| Sample Description | | Instrument | Agilent-6125B | | |
|---|---|---|---|---|---|
| Analyzed date: | 25-09-2020 | Probe: | ESI | Probe Bias: | +4.5kv |
| Analyst: | Dr.AR-SBio | Nebulizer Gas Flow: | 1.5L/min | Detector: | 1.5kv |
| Sample: | Pep-177 SGRGKQGGKARA | CDL: | -20.0v | T. Flow: | 0.2ml/min |
| M.W.: | 1172.29 | CDL Temp.: | 250 °C | B. Conc.: | 50%H2O/50%ACN |

**Figure S28:** MS spectra for the H2A unmodified (1-12) peptide.

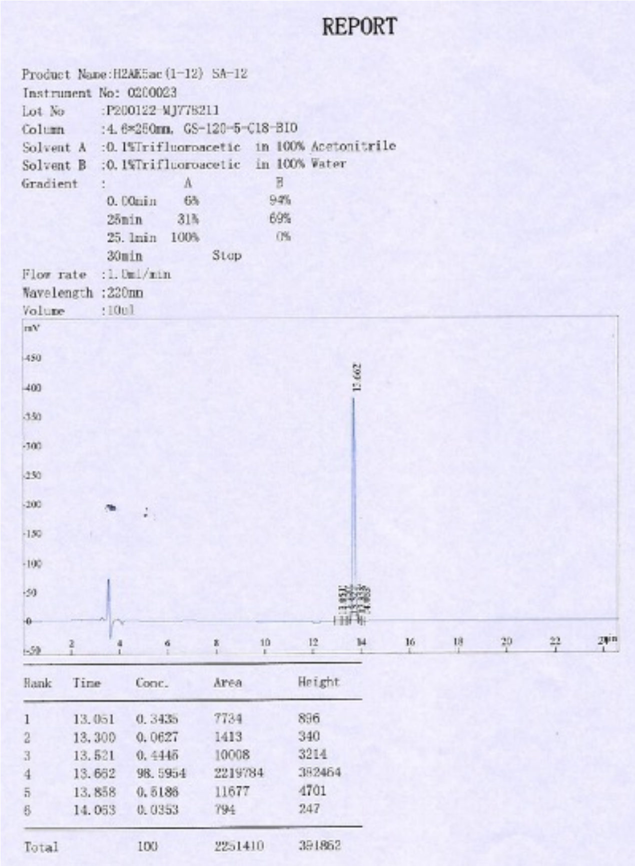

REPORT

Product Name:H2AK5ac(1-12) SA-12
Instrument No: 0200023
Lot No :P200122-MJ778211
Column :4.6*250mm, GS-120-5-C18-BIO
Solvent A :0.1%Trifluoroacetic in 100% Acetonitrile
Solvent B :0.1%Trifluoroacetic in 100% Water

| Gradient : | A | B |
|---|---|---|
| 0.00min | 6% | 94% |
| 25min | 31% | 69% |
| 25.1min | 100% | 0% |
| 30min | Stop | |

Flow rate :1.0ml/min
Wavelength :220nm
Volume :10ul

| Rank | Time | Conc. | Area | Height |
|---|---|---|---|---|
| 1 | 13.051 | 0.3435 | 7734 | 896 |
| 2 | 13.300 | 0.0627 | 1413 | 340 |
| 3 | 13.521 | 0.4445 | 10008 | 3214 |
| 4 | 13.662 | 98.5954 | 2219784 | 382464 |
| 5 | 13.858 | 0.5186 | 11677 | 4701 |
| 6 | 14.053 | 0.0353 | 794 | 247 |
| Total | | 100 | 2251410 | 391852 |

**Figure S29:** HPLC purity trace for the H2AK5ac (1-12) peptide.

MASS SPECTROMETRY REPORT

H2AK5AC (1-12)

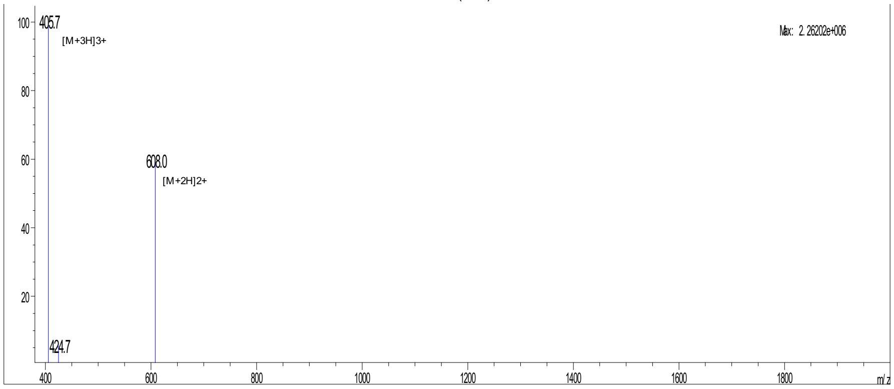


| | | | | | |
|---|---|---|---|---|---|
| Sample Description | | Instrument | Agilent-6125B | | |
| Analyzed date: | 2020-01-30 | Probe: | ESI | Probe Bias: | +4.5kv |
| Analyst: | YU | Nebulizer Gas Flow: | 1.5L/min | Detector: | 1.5kv |
| Sample: | H2AK5ac (1-12) SA-12 | CDL: | -20.0v | T. Flow: | 0.2ml/min |
| M.W.: | 1214.45 | CDL Temp.: | 250 °C | B. Conc.: | 50%H2O/50%ACN |
| Lot. No.: | P200122-MJ778211 | Block Temp.: | 200 °C | | |

**Figure S30:** MS spectra for the H2AK5ac (1-12) peptide.

**H3 unmodified (1-24)**

**HPLC REPORT**

| | | | | |
|---|---|---|---|---|
| Sample: | Pep-176 ARTKQTARKSTGGKAPRKQLATKA | | | Analyzed date: 01-10-2020 |
| Analyst: | Dr.RS-SBio | | | |
| Column: | Symmetrix ODS-R, 4.6*250mm, 5µm | | | |
| Solvent A | A: 0.1% Trifluoroacetic Acid in 100% Acetonitrile | | | |
| Solvent B | B: 0.1% Trifluoroacetic Acid in 100% Water | | | |
| Gradient: | | A | B | |
| | 0.0min | 1% | 99% | |
| | 25.0min | 26% | 74% | |
| | 25.1min | 100% | 0% | |
| | 30.0min | Stop | | |
| Volume: | 20µl | | | |
| Wavelength: | 220nm | | | |
| Flow rate: | 1.0ml/min | | | |

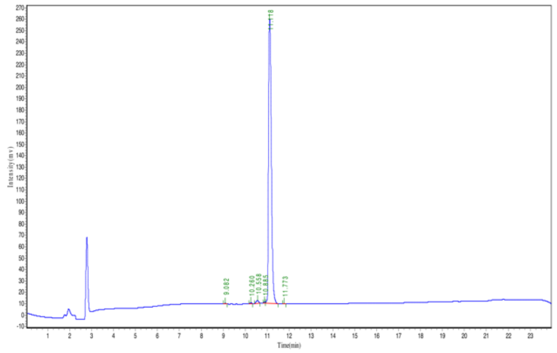

| Peak | Time | Height | Area | Conc. |
|---|---|---|---|---|
| 1 | 9.082 | 381.787 | 2113.503 | 0.0937 |
| 2 | 10.260 | 795.523 | 4257.644 | 0.1887 |
| 3 | 10.558 | 2550.401 | 18417.207 | 0.8164 |
| 4 | 10.885 | 1037.583 | 4749.035 | 0.2105 |
| 5 | 11.118 | 249803.641 | 2225080.250 | 98.6288 |
| 6 | 11.773 | 276.968 | 1395.949 | 0.0619 |
| Total | | | | 100.000 |

**Figure S31:** HPLC purity trace for the H3 unmodified (1-24) peptide.

**H3 unmodified (1-24)**

**MASS SPECTROMETRY REPORT**

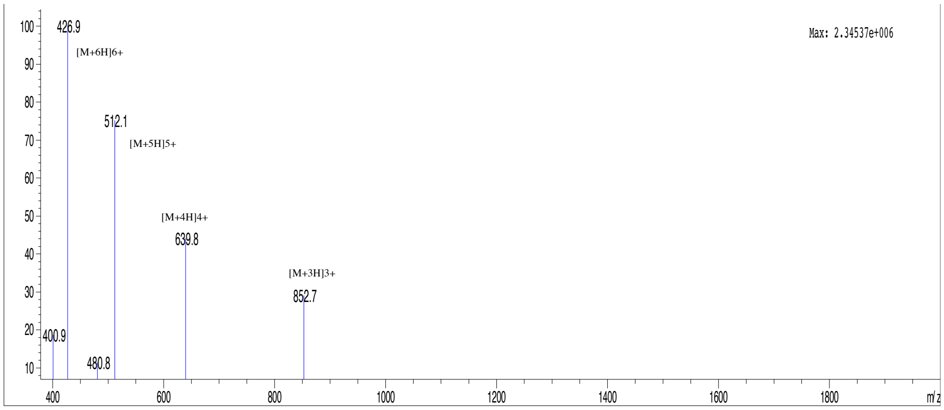


| | | | | | |
|---|---|---|---|---|---|
| Sample Description | | Instrument | Agilent-6125B | | |
| Analyzed date: | 02-10-2020 | Probe: | ESI | Probe Bias: | +4.5kv |
| Analyst: | Dr.RS-SBio | Nebulizer Gas Flow: | 1.5L/min | Detector: | 1.5kv |
| Sample: | Pep-176 ARTKQTARKSTGGKAPRKQLATKA | CDL: | -20.0v | T. Flow: | 0.2ml/min |
| M.W.: | 2554.94 | CDL Temp.: | 250 °C | B. Conc.: | 50%H2O/50%ACN |

**Figure S32:** MS spectra for the H3 unmodified (1-24) peptide.

**HPLC REPORT**
**H3K14ac (9-19)**

| | | |
|---|---|---|
| Sample: | Pep-174 KSTGGK(ac)APRKQ | Analyzed date: 21-09-2020 |
| Analyst: | Dr.RS-SBio | |
| Column: | Symmetrix ODS-R, 4.6*250mm, 5μm | |
| Solvent A | A: 0.1% Trifluoroacetic Acid in 100% Acetonitrile | |
| Solvent B | B: 0.1% Trifluoroacetic Acid in 100% Water | |

Gradient:

| | A | B |
|---|---|---|
| 0.0min | 1% | 99% |
| 25.0min | 26% | 74% |
| 25.1min | 100% | 0% |
| 30.0min | Stop | |

| | |
|---|---|
| Volume: | 20μl |
| Wavelength: | 220nm |
| Flow rate: | 1.0ml/min |

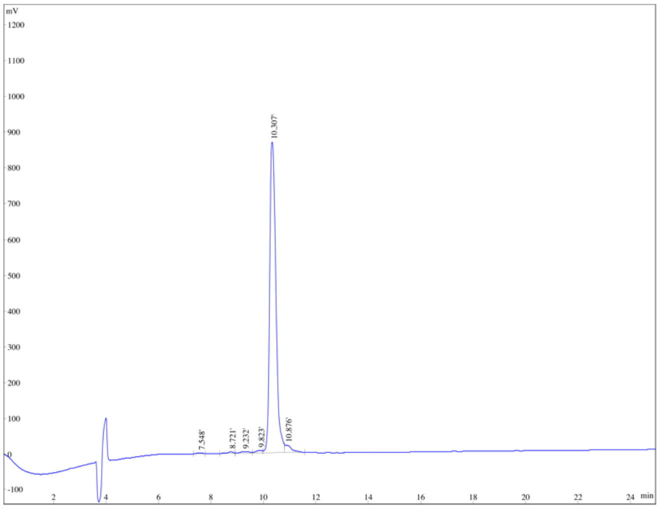


**Figure S33:** HPLC purity trace for the H3K14ac (9-19) peptide.

**H3K14ac (9-19)**
**MASS SPECTROMETRY REPORT**

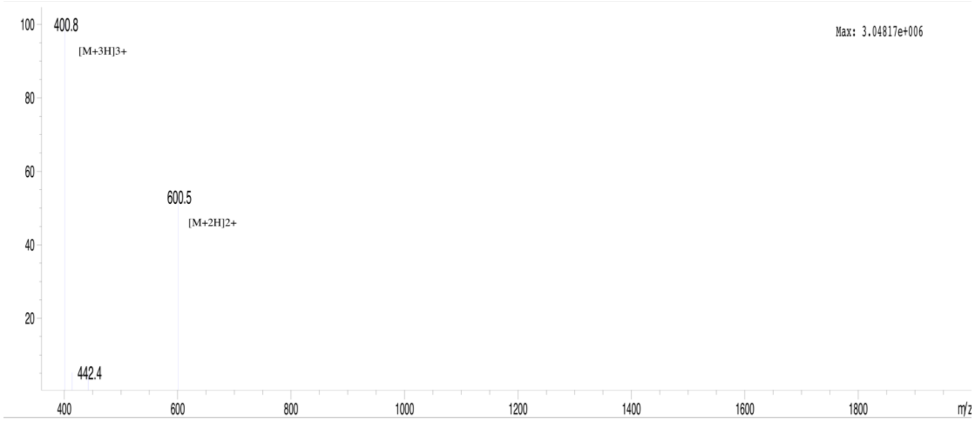


| | | | | | |
|---|---|---|---|---|---|
| Sample Description | | Instrument | Agilent-6125B | | |
| Analyzed date: | 21-09-2020 | Probe: | ESI | Probe Bias: | +4.5kv |
| Analyst: | Dr.AR-SBio | Nebulizer Gas Flow: | 1.5L/min | Detector: | 1.5kv |
| Sample: | Pep-174 KSTGGK(ac)APRKQ | CDL: | -20.0v | T. Flow: | 0.2ml/min |
| M.W.: | 1199.36 | CDL Temp.: | 250 °C | B. Conc.: | 50%H2O/50%ACN |

**Figure S34:** MS spectra for the H3K14ac (9-19) peptide.
